# Activity-dependent lateral inhibition signals for synaptic competition

**DOI:** 10.1101/625616

**Authors:** Satoshi Fujimoto, Marcus N. Leiwe, Shuhei Aihara, Richi Sakaguchi, Yuko Muroyama, Reiko Kobayakawa, Ko Kobayakawa, Tetsuichiro Saito, Takeshi Imai

**Affiliations:** Graduate School of Medical Sciences, Kyushu University, Fukuoka 812-8582, Japan; Laboratory for Sensory Circuit Formation, RIKEN Center for Developmental Biology, Kobe 650-0047, Japan; Graduate School of Biostudies, Kyoto University, Kyoto 606-8501, Japan; Department of Developmental Biology, Graduate School of Medicine, Chiba University, Chiba 260-8670, Japan; Institute of Biomedical Science, Kansai Medical University, Hirakata 573-1010, Japan; PRESTO and CREST, Japan Science and Technology Agency (JST), Saitama 332-0012, Japan

## Abstract

In developing brains, activity-dependent remodeling facilitates the formation of precise neuronal connectivity. Synaptic competition is known to facilitate synapse elimination; however, it has remained unknown how different synapses compete to each other within a postsynaptic cell. Here we investigate how a mitral cell in the olfactory bulb prunes all but one primary dendrite during the developmental remodeling process. We find that spontaneous activity generated within the olfactory bulb is essential. We show that strong glutamatergic inputs to one dendrite trigger branch-specific changes in RhoA activity to facilitate the pruning of the remaining dendrites: NMDAR-dependent local signals suppress RhoA to protect it from pruning; however, the subsequent neuronal depolarization induces neuron-wide activation of RhoA to prune non-protected dendrites. NMDAR-RhoA signals are also essential for the synaptic competition in the barrel cortex. Our results demonstrate a general principle whereby activity-dependent lateral inhibition across synapses establishes a discrete receptive field of a neuron.

## INTRODUCTION

In the mammalian nervous system, neurons initially form exuberant connections, and then undergo activity-dependent remodeling to form mature neuronal circuits. Compared to the neurite guidance process (Kolodkin and Tessier-Lavigne, 2011), the mechanisms of the remodeling process remains poorly understood. Hebbian plasticity explains how some synapses are strengthened (Hebb, 1949; Luo, 2021; Shatz, 1992); however, it is not fully understood how other synapses are actively eliminated (Lichtman and Colman, 2000; Wong and Ghosh, 2002). Synaptic competition is known to facilitate synapse elimination during the remodeling process. Inter-neuronal competition has been observed for axons (e.g., the neuromuscular junction and the climbing fiber – Purkinje cell synapses) (Lichtman and Colman, 2000; Watanabe and Kano, 2011): Initially multiple axons innervate a single synaptic target; however, all but one of these axons are pruned during development. Intra-neuronal competition is known for dendrites (e.g., Layer 4 neurons in the barrel cortex and mitral cells in the olfactory bulb): One or a few dendrites receiving a common input are stabilized, whereas the others are pruned within a neuron (Wong and Ghosh, 2002). In both cases, each postsynaptic cell selects just one group of presynaptic inputs as winners and eliminates surplus ones as losers, suggesting that activity-dependent events happening in a postsynaptic cell plays a key role in competition (Figure 1A). However, it has long been an enigma as to how a postsynaptic cell stabilizes just one winner and eliminates all the others as losers. It has been postulated that hypothetical “punishment signals” emitted from the winner synapses may facilitate elimination of losers (Lichtman and Colman, 2000). Yet, its exact nature remains entirely unknown. It was also puzzling how the hypothetical punishment signals solely act on losers, but not on winners.

**Figure 1.**
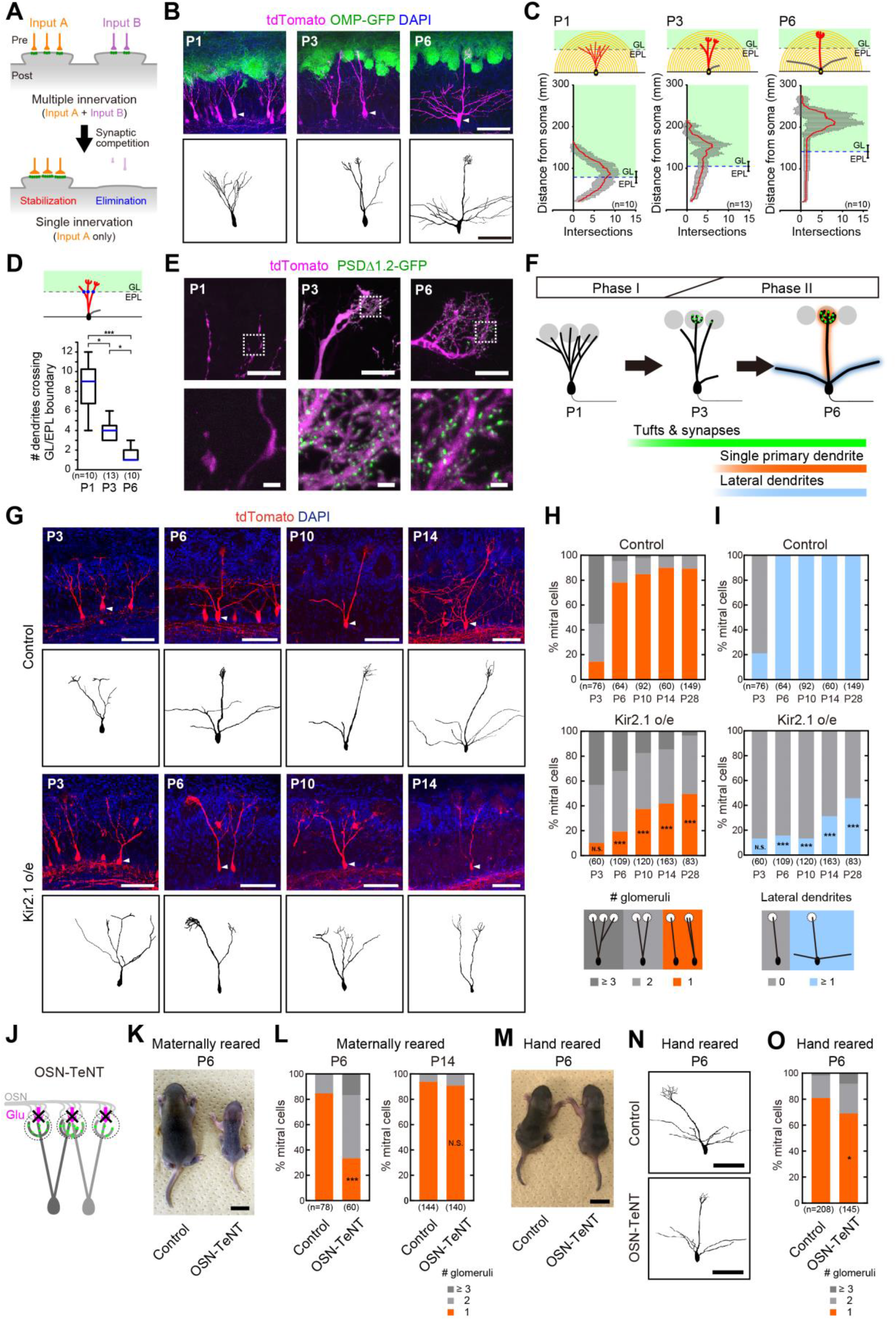
Neuronal activity in mitral cells is required to establish single primary dendrites. **(A)** A general scheme of synaptic competition during circuit remodeling. **(B)** Representative images of P1, P3, P6 mitral cells labelled with tdTomato (magenta). OMP-GFP knock-in mice were used to label OSN axons (green). Samples were cleared with SeeDB2. Representative neurons (white arrowheads) were reconstructed by Neurolucida and are shown below. A scale bar, 100 μm. **(C)** Modified Sholl analysis of mitral cell dendrites at P1, P3 and P6. Only primary dendrites extending to the glomerular layer (GL) were quantified. Data are the mean ± SD. EPL, external plexiform layer. **(D)** The number of dendrites extending into the GL at P1, P3 and P6. The number of dendrites crossing the GL/EPL boundary was quantified (blue line represents the median, the boxes are the inter-quartile range, and the whiskers represent the minimum and maximum). *p<0.05, ***p<0.001 (Kruskal-Wallis test and post hoc Dunn’s multiple comparison test). n, number of mitral cells. **(E)** Postsynaptic structures on primary dendrites at P1, P3 and P6. Primary dendrites (magenta) and PSDΔ1.2-GFP puncta (green) are shown. Scale bars, 20 μm (top) and 2 μm (bottom). **(F)** Schematic representation of mitral cell development. After the initial extension of neurites, synapses and tufts emerge heterogeneously on dendrites by P3, then pruning of dendrites occurs to leave a single primary dendrite by P6. **(G)** Kir2.1 overexpression in mitral cells perturbs the pruning of mitral cell dendrites and the formation of lateral dendrites when compared to the controls. Representative mitral cell morphologies at P3, P6, P10, and P14 and their reconstructions are shown. Scale bars, 100 μm. **(H)** Quantification of primary dendrite pruning in control (top) or Kir2.1-overexperssing (bottom) mitral cells. Percentages of mitral cells with a connection to single (1), double (2), or multiple (≥ 3) glomeruli at P3, P6, P10, P14, and P28 were analyzed. We did not distinguish between tuft(+) vs. tuft(−) dendrites in this quantification. N.S., non-significant, ***p<0.001 (χ^2^ test, vs. control). n, number of mitral cells. **(I)** Quantification of lateral dendrite formation. The percentages of mitral cells with/without lateral dendrites at P3, P6, P10, P14, and P28 are shown. N.S., non-significant, ***p<0.001 (χ^2^ test, vs. control). n, number of mitral cells. **(J)** Synaptic transmission from OSNs were blocked in OSN-TeNT mice. **(K)** Representative images of maternally reared control and OSN-TeNT mice at P6. A scale bar, 10 mm. **(L)** Quantification of dendrite pruning in maternally reared OSN-TeNT mice at P6 and P14. ***p<0.001 (χ^2^ test, vs. control). N.S., non-significant, (χ^2^ test, vs. control). n, number of mitral cells. **(M)** Representative images of hand reared control and OSN-TeNT mice at P6. A scale bar, 10 mm. **(N)** Representative dendritic reconstructions in hand reared control and OSN-TeNT mice. A scale bar, 100 μm. **(O)** Quantification of dendrite pruning in hand reared control and OSN-TeNT mice at P6. *p<0.05 (χ^2^ test, vs. control). n, number of mitral cells.

To study mechanisms of synaptic competition, mitral cells in the olfactory bulb can serve as an excellent model system. In each glomerulus of the olfactory bulb, sensory inputs from a specific type of olfactory sensory neurons (OSNs) are relayed to dendrites of 20-50 mitral and tufted cells. In the adult, a mitral cell typically has a single primary dendrite (also known as an apical dendrite) connecting to a single glomerulus, as well as several lateral dendrites (also known as basal dendrites) which solely receive inhibitory inputs from interneurons in the external plexiform layer. The discrete connectivity of primary dendrites to glomeruli is the basis for the receptive field of a mitral cell representing a specific type of odorant receptor (Imai, 2014). It has been known that this discrete connectivity is a result of the remodeling process (Malun and Brunjes, 1996; Santacana et al., 1992). However, it has remained unknown how a mitral cell establishes “just one” dendritic connection to a glomerulus (Imai, 2014). Revealing the underlying mechanisms will provide a fundamental principle in circuit remodeling in general.

## RESULTS

### Developmental remodeling of mitral cell dendrites

To understand the mechanisms of dendrite pruning in mitral cells, we took a genetic approach using *in utero* electroporation (Imamura and Greer, 2013; Muroyama et al., 2016; Saito, 2006). We electroporated plasmid DNA at E12 to label and manipulate a subset of mitral cells. The Cre-loxP system was used to sparsely label mitral cells (Figure 1B). Olfactory bulb samples were cleared, and volumetric fluorescence images were analyzed (Figures S1A and S1B; Movie S1).

During development, mitral cells initially extend multiple dendrites toward the glomerular layer (Blanchart et al., 2006; Lin et al., 2000; Malun and Brunjes, 1996; Matsutani and Yamamoto, 2000). However, by postnatal day 6 (P6) most mitral cells have a single primary dendrite with tufted structure extending to a single glomerulus (Figures 1C, 1D, S1C and S1D). Day-to-day quantification indicated that the pruning of surplus dendrites occurs between P4-5 (Figure S1D). We also examined the formation of glutamatergic synapses using PSDΔ1.2-GFP, which labels mature glutamatergic synapses (Arnold and Clapham, 1999; Hayashi-Takagi et al., 2015). Prominent PSDΔ1.2-GFP puncta began to appear in tufted structures from P3, just prior to the dendrite pruning (Figure 1E).

These observations indicate that the remodeling of mitral cell dendrites occurs in a stepwise manner (Figure 1F). During Phase I (≤P3), the number of dendrites is gradually reduced to ∼4 and a tufted structure with excitatory synapses begins to appear in one or a few of them. During Phase II (P3-6), just one dendrite is further strengthened to have a ramified tufted structure, while surplus ones are pruned. Several lateral dendrites are also formed during this stage.

### Neuronal activity in mitral cells is essential for dendrite pruning

Neuronal activity often plays an important role in neuronal remodeling in the mammalian brain (Wong and Ghosh, 2002). However, previous attempts to determine a role for neuronal activity in mitral cell remodeling have been unsuccessful (Lin *et al*., 2000; Ma et al., 2014; Matsutani and Yamamoto, 2000; Nishizumi et al., 2019). As all of these studies manipulated neuronal activity only in OSNs, we directly silenced neuronal activity in mitral cells by overexpressing a gene coding for the inward rectifying potassium channel, Kir2.1 (Figure S1E and S1F) (Burrone et al., 2002). Kir2.1 overexpression did not affect the initial growth and remodeling of dendrites during Phase I; however, during Phase II, dendrite pruning was significantly perturbed (Figures 1G, 1H, S1G and S1K). By P6, 78.1% of control mitral cells established connection to just one glomerulus; however, only 19.3% of Kir2.1-expressing mitral cells established connection to a single glomerulus at this stage (Figure 1H). The formation of lateral dendrites was also precluded (Figures 1I and S1G), while axons were not affected at P6 (Figure S1H). Tufted structures and glutamatergic synapses were formed in the majority of primary dendrites of Kir2.1-expressing mitral cells, suggesting that neuronal activity is required for the pruning of surplus dendrites that can otherwise form mature glutamatergic synapses (Figure S1I). A non-conducting mutant of Kir2.1 (Burrone *et al*., 2002) did not affect dendrite pruning (Figure S1J).

We examined whether neuronal activity in OSNs is required. Firstly, we confirmed that a unilateral naris occlusion from P0 caused no change in primary dendrite pruning at P6, excluding a role for sensory-evoked activity (Figure S2A). We next examined the role for any kinds of activity derived from OSNs. We generated a mouse line expressing tetanus toxin light chain (TeNT) in OSNs (*OMP-Cre;R26-CAG-LoxP-TeNT*, designated OSN-TeNT); TeNT cleaves VAMP2 and thus inhibits synaptic vesicle release (Figure 1J, S2D and S2G-S2I) (Yamamoto et al., 2003; Yu et al., 2004). OSN-TeNT mice showed delay in dendrite pruning process (Figure 1L) as well as growth retardation (Figure 1K and S2E); the growth delay is likely due to anosmia-related suckling defects (Figures S2G-S2I). We obtained similar results for the OSN-specific CNGA2 mutant (*OMP-Cre; Cnga2*^*fl/fl*^) (Figure S2B, S2C).

We, therefore, tried to rescue the growth retardation by hand rearing new-born animals. We fed powdered milk to OSN-TeNT mice manually every 2 hours from P1-6 in a warm chamber (Leiwe et al., 2020). This hand rearing partially rescued the growth delay (Figure 1M), and these OSN-TeNT mice were of a similar weight (98.9 ± 11.1%, mean ± SD) to hand-reared controls, and 62.2 ± 12.9% (mean ± SD) of maternally reared controls (Figure S2F). The hand-rearing also largely rescued the defective dendrite pruning seen in mother-reared OSN-TeNT animals (Figure 1N). Now, the difference between hand-reared control and OSN-TeNT mice was small, if any, at P6 (Figure 1O). Thus, neuronal activity derived from OSNs plays a minimal role, if any, for the activity-dependent pruning of mitral cell dendrites.

### Spontaneous activity in awake neonatal mice

Our genetic experiments raised the possibility that spontaneous activity in mitral cells is important for dendrite pruning. Spontaneous activity is known to occur in other sensory systems during development (Ackman et al., 2012; Babola et al., 2018; Feller et al., 1996; Mizuno et al., 2018; Tritsch et al., 2007), but it has not been investigated in the olfactory bulb. We performed *in vivo* two-photon Ca^2+^ imaging of the olfactory bulb in awake newborn mice using mitral/tufted cell-specific GCaMP3 and GCaMP6f mouse lines (transgenic *Pcdh21-Cre;Ai38* and *Thy1-GCaMP6f*, designated M/T-GCaMP3/6f hereafter) (Dana et al., 2014; Iwata et al., 2017; Zariwala et al., 2012) (Figure 2A). Awake neonatal mice at P4 demonstrated spontaneous neuronal activity in the glomerular layer of the olfactory bulb (Figure 2B). Spontaneous activity was not seen in anesthetized pups (Figures S3A and S3B), which was similar to earlier studies in other sensory systems (Ackman *et al*., 2012; Babola *et al*., 2018; Mizuno *et al*., 2018). We did not observe any patterned spontaneous activity in OSN axons in OSN-specific GCaMP3 mice (*OMP-tTA;TRE-GCaMP3*) (Figure 2C). Furthermore, robust spontaneous activity in mitral/tufted cells was observed even after naris occlusion (Figure 2D) and in OSN-TeNT mice (Figure S2J). Thus, the spontaneous activity in the olfactory bulb can be generated without synaptic inputs from OSNs.

**Figure 2.**
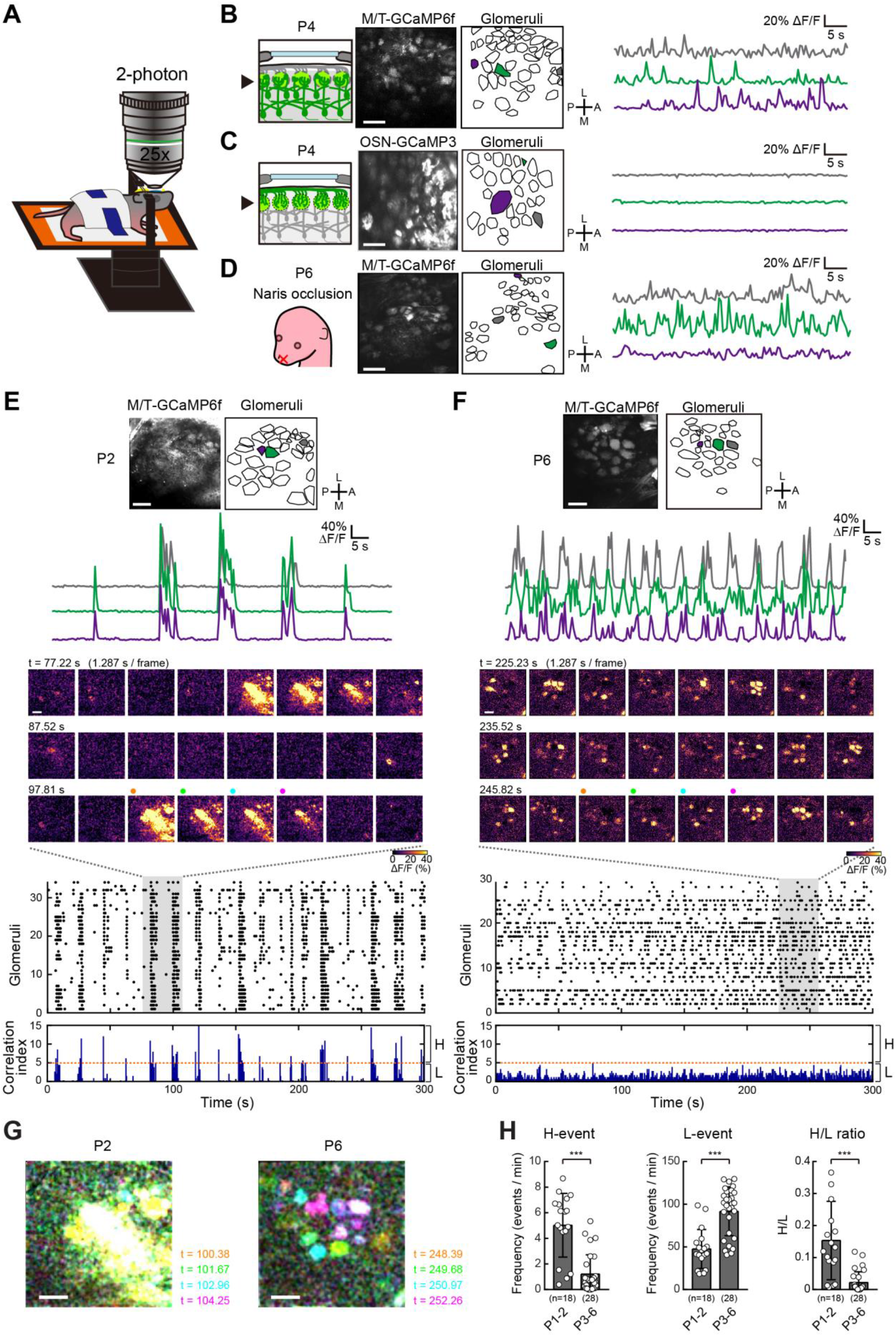
*In vivo* imaging of spontaneous activity in awake neonates. **(A)** Schematic representation of the imaging set up. A cranial window was made on the left olfactory bulb, and dental cement was used to attach a metal bar for head fixing. **(B)** GCaMP6f was specifically expressed in mitral/tufted cells and glomerular layer was imaged (left). Representative traces are shown (right) for 3 glomeruli (shown in gray, green, and purple), displaying spontaneous activity in awake animals. We obtained similar results with M/T-GCaMP3 (data not shown). A, anterior; P, posterior; M, medial; L, lateral.. **(C)** OSN-GCaMP3 did not show patterned spontaneous activity in any glomeruli. **(D)** Naris occlusion from P0-P6 does not affect spontaneous activity occurring in mitral cells. **(E, F)** Representative P2 (**E**) and P6 (**F**) imaging sessions (top), with individual traces shown below (3 glomeruli). A montage of serial ΔF/F images (middle, highlighted in gray area in the raster plot), raster plots, and correlation index histograms (bottom, See Methods) are also shown. A correlation index threshold of 5 (orange dotted lines) was selected to differentiate between highly correlated (H-) and lowly correlated (L-) events. See also Movie S2, S3. **(G)** Color representation of P2 and P6 spontaneous activity. Four consecutive images marked in **E** and **F** are displayed in four different colors. **(H)** Developmental changes in H-event frequency, L-event frequency, and the ratio of H to L events. The data was grouped into two phases (P1-2 and P3-6) for statistical analysis. As the H event frequency and H/L ratio were not normally distributed, Mann Whitney U tests were performed. For the L event frequency, a two-tailed t-test was performed. *p<0.05, **p<0.01, ***p<0.001. n, number of imaging sessions. All error bars represent SD. Scale bars, 100 μm. More detailed data for P1-P6 are shown in Figure S3.

We observed that the pattern of spontaneous activity changes during development. Spontaneous activity was synchronized among many glomeruli within 100-150 μm at P1-2, but became de-synchronized afterwards Figures 2E, 2F,2G, S3C and S3F; Movies S2 and S3). Based on the correlation index, each spike was classified into either highly correlated (H-) events and more sporadic lowly correlated (L-) events (See Methods) (Siegel et al., 2012). A high ratio of H-events was observed only at P1-2, and L-events dominated afterward (Figures 2H and S3G). At all stages, the activity was synchronized within each glomerulus (Maher et al., 2009; Schoppa and Westbrook, 2001). The spike frequency and amplitude were variable among glomeruli and ages (Figures S3D and S3E).

### Glutamatergic neurotransmission among mitral/tufted cells is critical

To examine whether the spontaneous activity is generated intrinsically in the olfactory bulb, we performed two-photon Ca^2+^ imaging of acute olfactory bulb slices *ex vivo*, thus removing any external inputs (Figure 3A). Similar age-specific patterns of spontaneous activity were observed in these preparations as *in vivo* (Figures 3B and 3C; Movies S4 and S5). H-events were more frequent at P2 than at P6 (Figure S4A), and a high correlation between glomeruli was found for many pairs at P2, but not at P6 (Figure S4B). At P2, the H-events were in fact propagating waves (Figures S4C-J). Thus, the age-specific patterns of spontaneous activity are generated within the OB circuits.

**Figure 3.**
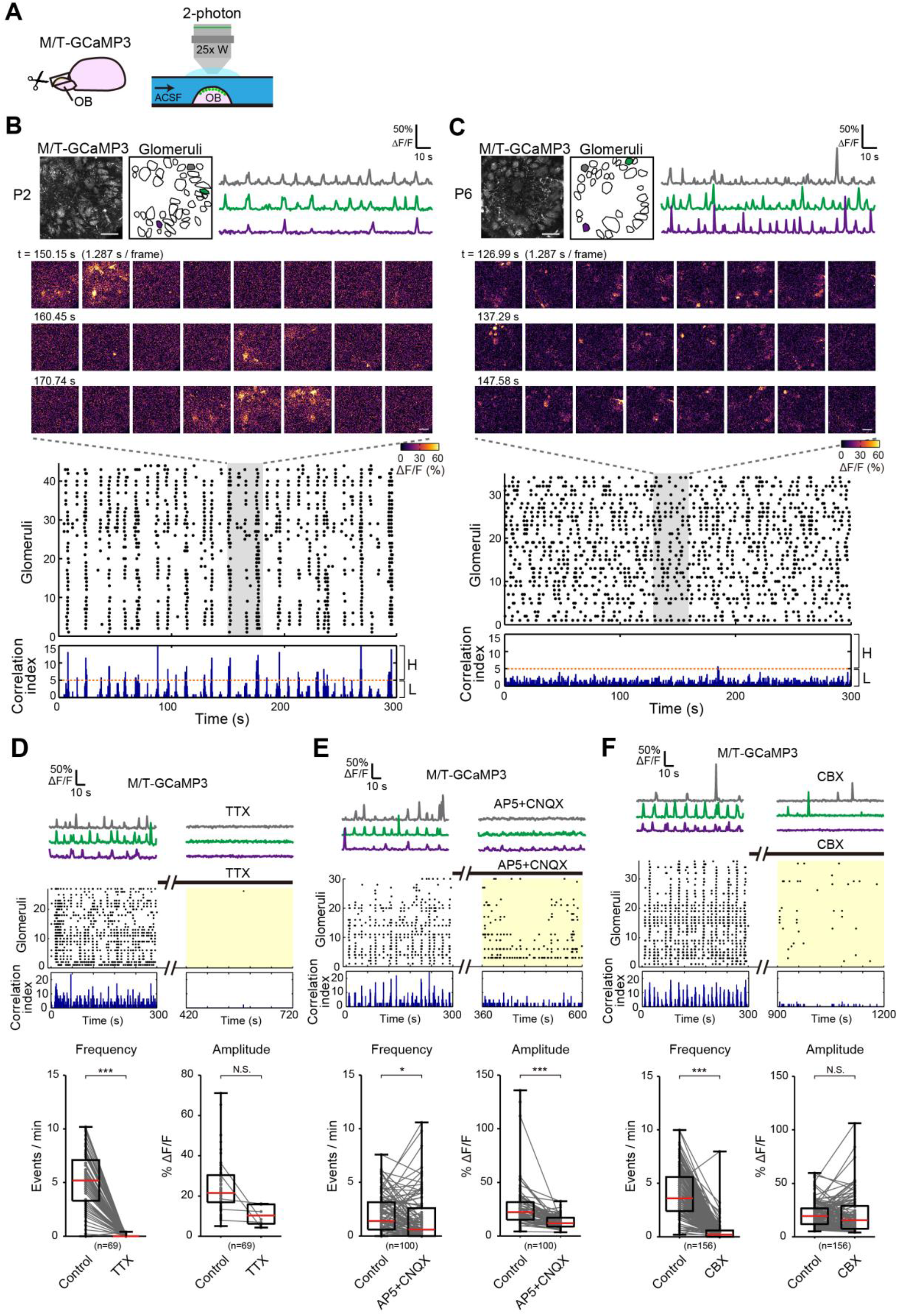
Spontaneous activity in the isolated olfactory bulb *ex vivo*. **(A**) Schematic of the slice imaging set up. Isolated olfactory bulbs (OB) were continuously perfused with artificial cerebrospinal fluid (ACSF). **(B, C)** Olfactory bulb slices of M/T-GCaMP3 mice were imaged at P2 (**B**) and P6 (**C**). Representative imaging sessions (top-left), with individual traces shown (top-right, shown in gray, green, and purple). A montage of serial ΔF/F images (middle, highlighted in shadowed area in the raster plot), raster plots, and correlation index histograms (bottom) are also shown. A correlation index threshold of 5 (orange dotted lines) was selected to differentiate between highly correlated (H-) and lowly correlated (L-) events. We obtained similar results with M/T-GCaMP6f. Scale bars, 100 μm. See also Movies S4 and S5. **(D)** Both the frequency and amplitude of glomerular spikes were reduced when TTX was added to P2 olfactory bulb slices. n = 69 glomeruli from 3 animals. N.S., non-significant, ***p<0.001 (Wilcoxon signed-rank test). **(E)** Application of AP5 and CNQX shows that ionotropic glutamate receptors are required for spontaneous activity in P2-3 olfactory bulb slices. n = 100 glomeruli from 4 animals. *p<0.05, ***p<0.001 (Wilcoxon signed-rank test). **(F)** Application of Carbenoxolone (CBX) shows that gap junctions are also required for spontaneous activity in P2-3 olfactory bulb slices. n = 156 glomeruli from 4 animals. N.S., non-significant, ***p<0.001 (Wilcoxon signed-rank test).

We investigated the possible origin of spontaneous activity using pharmacological manipulations of isolated olfactory bulbs *ex vivo*. Spontaneous activity was mostly eliminated by tetrodotoxin (TTX), a sodium channel blocker (Figure 3D). When we applied inhibitors to NMDA- and AMPA-type ionotropic glutamate receptors (AP5 and CNQX, respectively), both the amplitude and frequency of spontaneous activity were reduced (Figures 3E and S5A). Inhibitors for gap junctions also reduced the frequency of spontaneous activity (Figures 3F and S5B). GABA is unlikely to be involved: GABA was inhibitory toward mitral cells already at P2 (Figure S5C); moreover, we did not find any defects in dendrite pruning in mice deficient for NKCC1, which is essential for excitatory action of GABA (Figure S5D) (Wang and Kriegstein, 2008). Thus, glutamatergic inputs and gap junctions are important.

We therefore investigated the source of glutamatergic synaptic inputs. In the isolated olfactory bulb, mitral/tufted cells are the only possible source of glutamatergic inputs. Mitral/tufted cells are known to release glutamate not only from axons, but also from dendrites (Isaacson, 1999; Schoppa and Westbrook, 2002; Urban and Sakmann, 2002). We therefore tested the possibility that glutamatergic transmission among mitral/tufted cells is required for the dendrite pruning. We generated *Pcdh21-Cre;R26-CAG-loxP-TeNT* (designated M/T-TeNT) mice, in which TeNT is specifically expressed in mitral/tufted cells (Figures 4A and S6A). Unlike OSN-TeNT mice, M/T-TeNT mice did not show growth defects (Figure S6B). As we used the transgenic *Pcdh21-Cre* line, a small fraction of mitral/tufted cells may have failed to express Cre (Iwata *et al*., 2017). Nevertheless, the frequency and amplitude of spontaneous activity was reduced in M/T-TeNT mice *in vivo* (Figures 4B, 4C, and S6C) and *ex vivo* (Figures S6D-S6F). Morphological analysis revealed that M/T-TeNT mice show a significant reduction in the percentage of mitral cells innervating a single glomerulus, at both P6 and P14 (Figures 4D and 4E). Thus, glutamatergic neurotransmission among mitral/tufted cells is a major source of the spontaneous activity and is essential for the dendrite pruning of mitral cells (Figure 4F).

**Figure 4.**
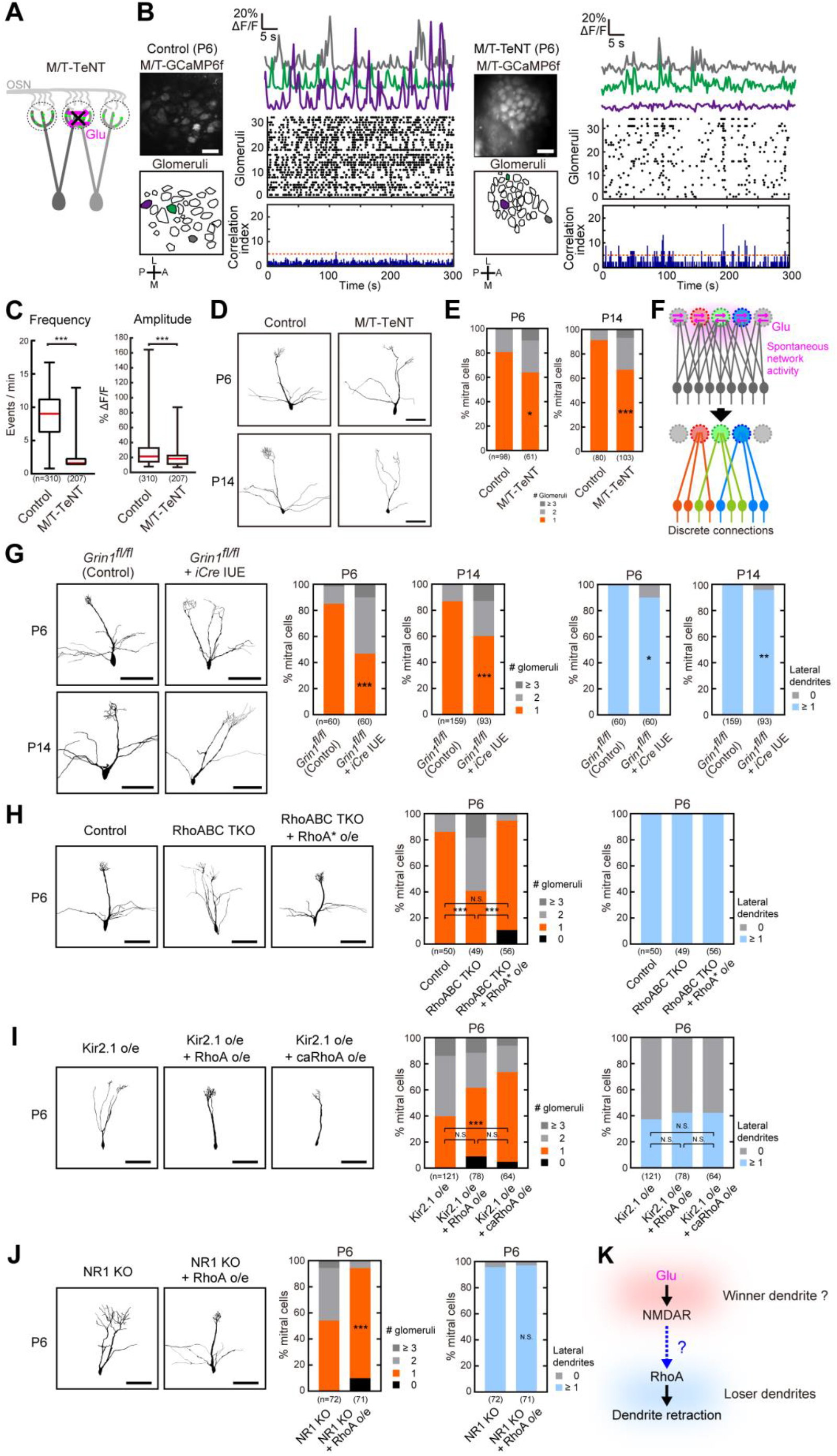
Glutamatergic inputs facilitate dendrite pruning via NMDARs and RhoA. **(A)** Neurotransmissions among mitral/tufted cells were blocked in M/T-TeNT mice. **(B)** Crossing M/T-TeNT mice with M/T-GCaMP6f mice allowed us to visualize neuronal activity in M/T-TeNT mice at P6. Representative Ca^2+^ traces and their raster plots are shown, along with their correlation index histograms (littermate control, left; mutant, right). Scale bars, 100 μm. **(C)** Boxplots showing the changes in glomerular firing frequency and amplitude. The frequency of glomerular spikes was drastically reduced (***p<0.001, Mann-Whitney U test) while the amplitude was also decreased (*** p<0.001, Mann-Whitney U test). Data are from a total imaging time of 598.88 and 571.64 minutes, from 310 and 207 glomeruli, from 10 and 8 imaging sessions from 2 and 3 animals for littermate control and M/T-TeNT mice, respectively. **(D)** Representative dendritic reconstructions in control and M/T-TeNT mice at P6 and P14. Scale bars, 100 μm. **(E)** Quantitative analysis of dendrite pruning in control and M/T-TeNT mice at P6 and P14. *p<0.05, ***p<0.001 (χ^2^-test vs control). In the box plots, the red line represents the median, the box is the inter-quartile range, while the whiskers represent the extremes. **(F)** Summary schema, describing the nature of spontaneous activity in the OB, which is propagated along the M/T cell network via dendro-dendritic transmission. See also Figures S6A-S6F. **(G)** Conditional knockout of NR1. Cre plasmids were introduced into mitral cells of floxed-*Grin1* mice. Representative dendritic morphology, and the quantification of primary and lateral dendrites at P6 and P14 are shown. Sparse and global knockout experiments are shown in Figures S6G-S6I. Scale bars, 100 μm. **(H)** Triple knockout of RhoA, B, and C using CRISPR-Cas9 and *in utero* electroporation. The defective phenotype was rescued by a CRISPR-resistant RhoA plasmid (RhoA*). **(I)** Wild-type and a constitutively-active mutant of RhoA was overexpressed together with Kir2.1, demonstrating partial rescue. **(J)** RhoA was overexpressed in CRISPR-Cas9-mediated NR1-knockout neurons. Dendrite pruning was rescued by RhoA. Percentages of mitral cells with a connection to zero (0), single (1), double (2), or multiple (≥ 3) glomeruli were analyzed. ***p<0.001, *p<0.05, **p<0.01, N.S., non-significant (χ^2^-test vs control in **A**, χ^2^-test with Bonferroni correction in (B) and (C), χ^2^-test vs NR1 KO in (D)). n, number of mitral cells. Scale bars, 100 μm in (A)-(D). **(K)** A summary schema for genetic experiments. NMDARs should activate RhoA but only in remote dendrites.

### NMDARs facilitate dendrite pruning via RhoA

We therefore examined a possible role for NMDARs in dendrite pruning. We performed a sparse knockout of *Grin1* coding for NR1, an essential subunit of NMDARs. We found that dendrite pruning was impaired in NMDAR-deficient mitral cells at both P6 and P14 (Figures 4G and S6G). Sparse *Grin1* expression under a mitral/tufted cell-specific global *Grin1* knockout rescued the defective phenotype (Figure S6H-S6J). Thus, NMDARs regulate the pruning of mitral cell dendrites in a cell-autonomous fashion.

We next examined how NMDARs regulate dendrite pruning. Previous studies indicated that various types of Rho GTPases regulate activity-dependent dendrite morphogenesis, including growth and retraction (Li et al., 2002; Luo, 2000; Murakoshi et al., 2011; Sin et al., 2002). Using CRISPR-Cas9-based screening in mitral cells (Aihara et al., 2021), we found that Rho subfamily GTPases are required for dendrite pruning (Figure 4H). When all of the Rho subfamily GTPases (RhoA, RhoB, and RhoC) were knocked out in mitral cells (RhoABC TKO), dendrite pruning was prevented. Conversely, when RhoA was overexpressed in NMDAR-deficient mitral cells, the defective phenotype was largely restored (Figure 4J). We obtained similar results for the rescue of Kir2.1 overexpression (Figure 4I). Thus, RhoA mediates NMDAR-dependent dendrite pruning.

### Glutamatergic inputs via NMDARs triggers local suppression and global activation of RhoA

As glutamatergic synapses are predominantly localized at presumptive “winner” primary dendrites with tufted structures (Figure 1E), a puzzling question here is how NMDAR-RhoA signals selectively prune remotely-located weaker dendrites with less or no NMDARs (Figure 4K). To gain a mechanistic insight into the selectivity of the dendrite pruning, we examined the spatial patterns of RhoA activity upon NMDA bath application using a FRET biosensor for RhoA at P3 *ex vivo*. When we applied NMDA and Glycine to activate NMDARs in the presence of tetrodotoxin (TTX), RhoA was activated in the somata as well as many parts of the dendrites of mitral cells. Paradoxically, however, RhoA activity was decreased within the tufted structures of primary dendrites where glutamatergic synapses are highly enriched (Figures 5A-5C and S7A). The response polarity in tufts vs soma was unchanged under different doses of NMDA stimuli (Figures 5G and S7D), arguing against the possibility that different amounts of NMDAR signals produced the opposite response polarities. In contrast, High K^+^ stimulus increased RhoA activity at both somata and dendritic tufts (Figures 5A-5C). A mutant FRET sensor did not show any responses (Figures 5C). These results suggest that the sources of the stimuli are important: NMDAR inputs at tufted dendrites locally suppress RhoA activity and protect them from pruning; however, subsequent depolarization signals activate RhoA on a neuron-wide scale to facilitate the pruning of non-protected dendrites.

**Figure 5.**
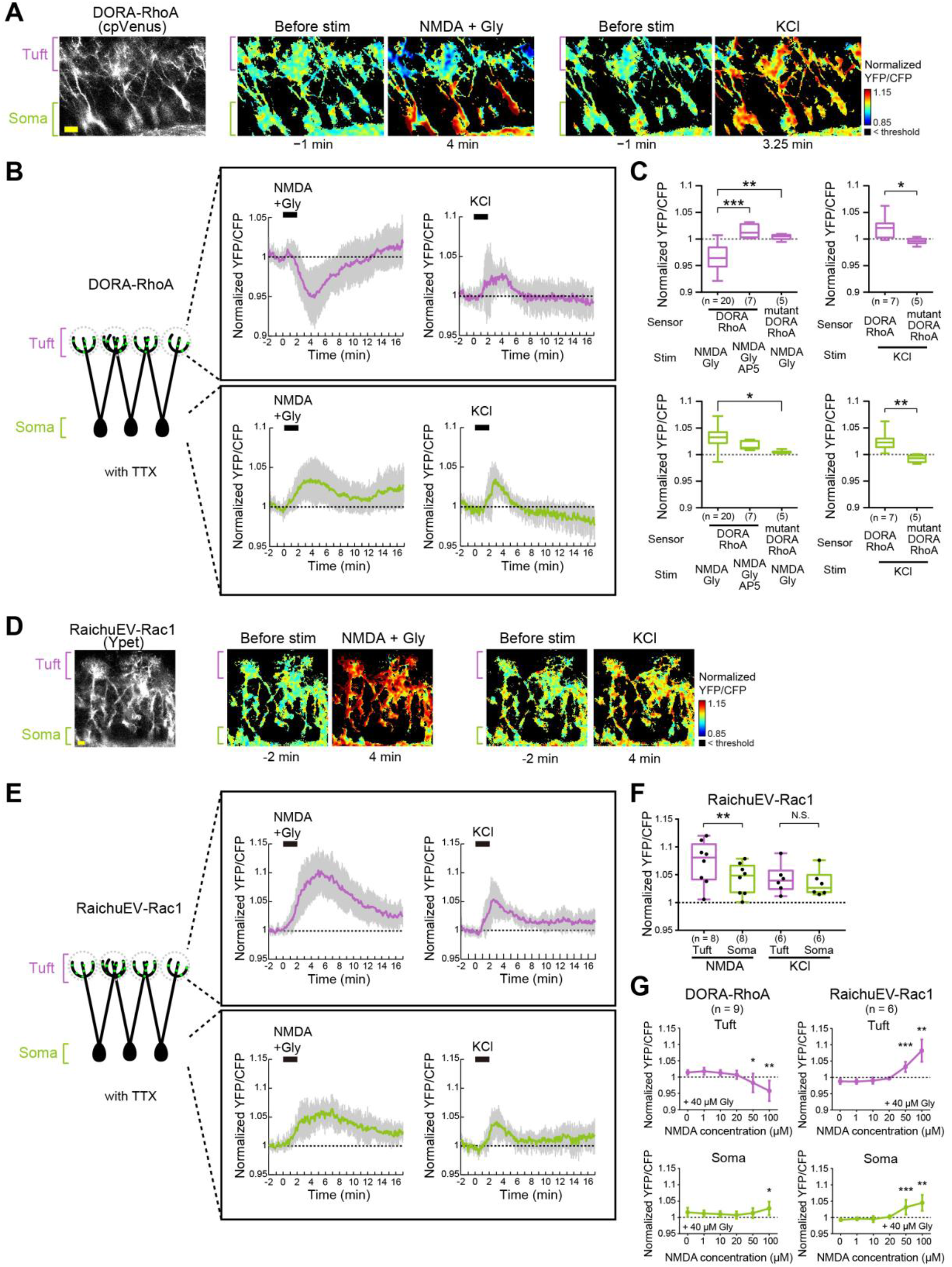
NMDAR stimulation induces RhoA suppression and Rac1 activation at the tuft, whereas depolarization induces a neuron-wide activation of RhoA. **(A)** FRET imaging of RhoA activity in mitral cells (P3 OB) with the DORA-RhoA biosensor under TTX. A cpVenus (YFP) image shows the morphology of mitral cells (left). Normalized YFP/CFP ratios before and after NMDAR stimulation (NMDA + Gly) and high K^+^ are shown. **(B)** Temporal FRET changes in RhoA activity at the tuft (top) and the soma (bottom) regions. **(C)** Quantitative analyses of RhoA activity. NMDAR antagonist AP5 and mutant DORA RhoA were used as negative controls. Mean values during 2-4 min are shown. ***p<0.001, **p<0.01, *p<0.05, (one-way ANOVA with Tukey post hoc tests in the left and t-tests in the right). **(D)** FRET imaging of Rac1 activity with the RaichuEV-Rac1 biosensor under TTX. A Ypet (YFP) image shows the morphology of mitral cells (left). Normalized YFP/CFP ratio before and after NMDAR stimulation and high K^+^ are shown. **(E)** Temporal FRET changes in Rac1 activity at the tuft (top) and the soma (bottom) regions. **(F)** Quantification of Rac1 activity at the tuft and the soma regions after NMDAR or K^+^ stimulation. Mean values during 2-4 min are shown. **p<0.01, N.S., non-significant (Paired t-tests) Scale bars, 20 μm. **(G)** Quantification of RhoA and Rac1 activity after NMDAR stimulation in different concentrations of NMDA. Mean values during 2-4 min are shown. *p<0.05, **p<0.01, ***p<0.001, (Paired t-tests between 0 and each concentration).

These results were in contrast to another Rho GTPase, Rac1, which is known to stabilize primary dendrites via actin fiber formation (Aihara *et al*., 2021): Upon NMDA application, Rac1 was highly activated in dendritic tufts but only weakly at somata (Figures 5D-5F). Together, these results suggest that NMDAR-dependent local signals at postsynapses induce RhoA suppression and Rac1 activation, which together protect them from pruning, whereas depolarization signals alone lead to RhoA activation to facilitate the dendrite pruning on a neuron-wide scale.

### Branch-specific dynamics of RhoA suggests lateral inhibition within a neuron

To directly prove this possibility, we next examined whether NMDAR inputs to one dendrite induce branch-specific changes in RhoA activity within the mitral cell *ex vivo*. We introduced the RhoA FRET sensor sparsely to perform FRET imaging of mitral cells at a single-cell resolution. Unfortunately, NMDA uncaging at one dendritic tuft was not possible, as it caused photobleaching of the FRET sensor. Instead, we performed NMDA bath application and focused on a group of mitral cells which have a thick-tufted primary dendrite (tuft(+)) and a non-tufted primary dendrite (tuft(−)). In these mitral cells, glutamatergic synapses are rich in tuft(+) but sparse in tuft(−) dendrites, allowing for selective activation NMDARs on tuft(+) dendrites (Figures S7B and S7C). NMDA application to this type of neurons indeed demonstrated branch-specific responses: RhoA activity tended to show a reduction at tuft(+) dendrites but an elevation at tuft(−) dendrites (Figures 6A,6B, 6E and 6G). Thus, NMDARs suppress RhoA activity in the vicinity of activated synapses, while activating RhoA in remote dendrites.

**Figure 6.**
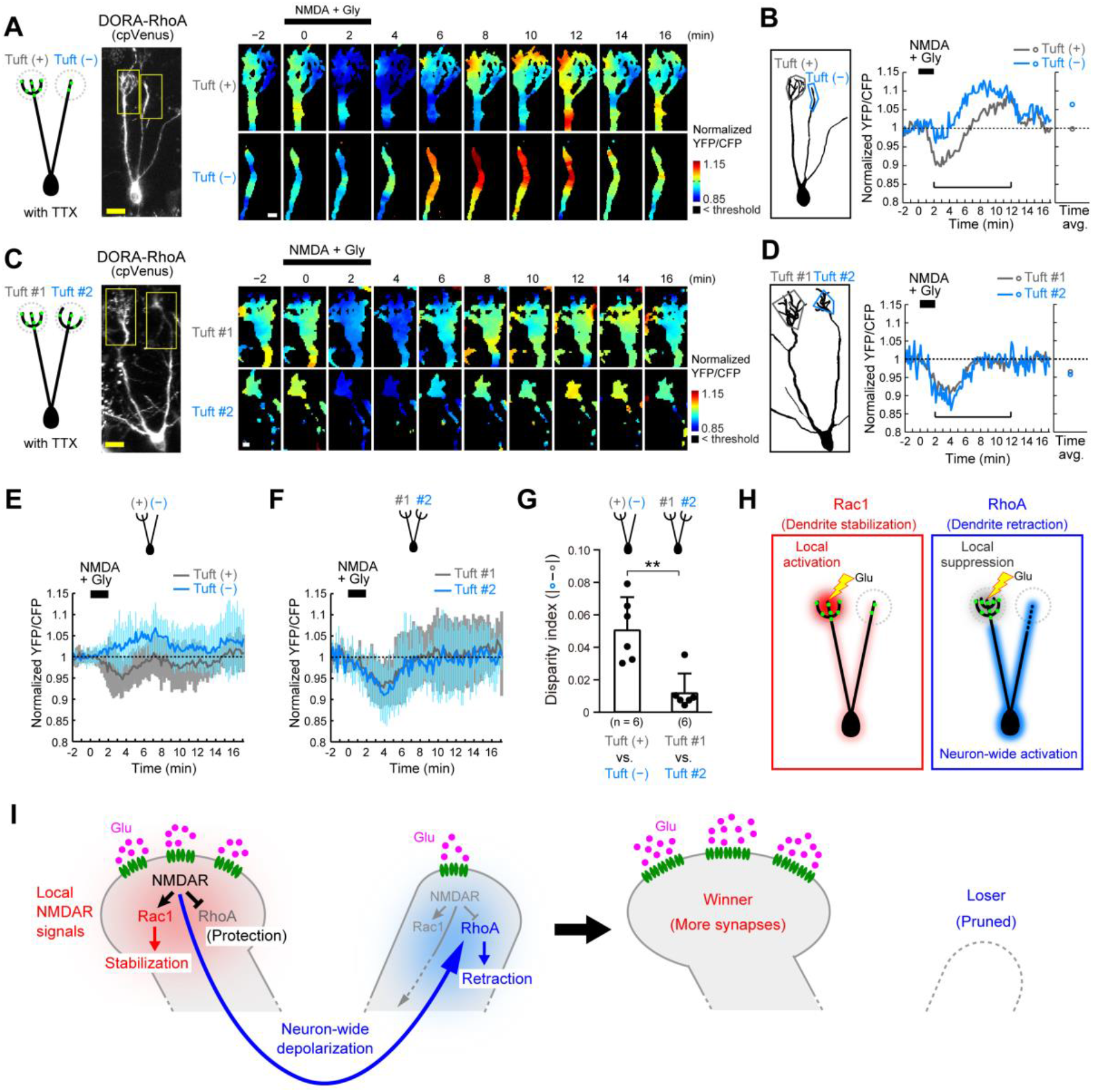
Branch-specific dynamics of RhoA by NMDAR inputs. **(A)** Single-cell analysis of RhoA activity upon NMDAR stimulation (age, P3). A representative mitral cell with a tuft(+) and a tuft(−) primary dendrites are shown. **(B)** Quantification of RhoA activity in the tuft(+) (gray) and the tuft(−) (blue) dendrites. Average values during 2-12 min are shown on the right. **(C)** A representative mitral cell with two tuft (+) primary dendrites (tuft #1 and tuft #2) are shown. **(D)** Quantification of RhoA activity in the two dendrites (tuft #1 in gray and tuft #2 in blue). Average values during 2-12 min are shown on the right. **(E)** Temporal changes in RhoA activity in tuft (+) and a tuft (−) primary dendrites. Data are the mean ± SD from 6 mitral cells. **(F)** Temporal changes in RhoA activity in tuft #1 and a tuft #2 primary dendrites. **(G)** Branch-specificity of RhoA responses were compared between mitral cells with one tuft(+) and one tuft(−) dendrite vs. those with two tuft(+) dendrites. The disparity index shows the difference in normalized YFP/CFP ratios between branches (mean values during 2-12 min). **p<0.01, Mann-Whitney U test **(H)** Schematic summary of FRET imaging experiments. **(I)** Our proposed model on the activity-dependent synaptic competition. Strong glutamatergic inputs at a presumptive “winner” dendrite leads to the local suppression of RhoA to protect it from pruning. The subsequent depolarization induces the neuron-wide activation of RhoA, leading to the pruning of non-protected dendrites (lateral inhibition). NMDAR-dependent local signals also activate Rac1 to stabilize the dendrite (Aihara *et al*., 2021). Scale bars, 20 μm in (A) and (C) left; 5 μm in (C) right.

We also examined another group of mitral cells with two tuft(+) primary dendrites. In these neurons, NMDA application showed RhoA suppression in both of the tuft(+) dendrites. The disparity of RhoA responses between dendrites was smaller for mitral cells with two tuft(+) dendrites (Figures 6C, 6D, 6F and 6G). These results indicate that the suppression vs. activation of RhoA reflects the relative weight of NMDAR-dependent local signals vs. depolarization signals representing the total synaptic inputs in the neuron. If only one of the dendrites receive strong glutamatergic inputs within several minutes of time window, the activated dendrite suppresses RhoA locally while remotely activating RhoA at the other dendrites. If two dendrites are strongly activated at the same timing, both of them will be protected (Figure 6D and F). Thus, RhoA acts as the “lateral inhibition” signals to facilitate the selective elimination of dendrites with weaker inputs.

### NMDAR-RhoA signals are also critical for the barrel circuit remodeling

Lastly, we examined whether lateral inhibition via NMDAR-RhoA signals is a general mechanism for the synaptic competition. It has been known that NMDARs are critical for synaptic competition in thalamocortical circuits. For example, layer 4 neurons in the barrel cortex are known to establish whisker-specific dendritic orientations through an NMDAR-dependent remodeling process (Espinosa et al., 2009; Mizuno et al., 2014). Here we found that the triple knockout of Rho subfamily GTPases (RhoA, B, C) abolishes dendritic orientation, similar to the NR1 knockout (Figure 7A-E). In the wild-type control, majority of dendrites were found within a major hollow, receiving inputs from one whisker; however, more dendrites were found within the septa and another hollow in the knockout neurons. We also found that defective dendrite pruning in NR1-deficient neurons, but not correct dendritic orientation, was rescued by RhoA overexpression. It has been reported that whisker-specific spontaneous activity is happening during the remodeling process (Mizuno *et al*., 2018). Thus, our results indicate that NMDAR-dependent RhoA signaling is critical for the activity-dependent elimination of L4 dendrites which fail to receive strong and correlated neuronal activity (Figure S7H).

**Figure 7.**
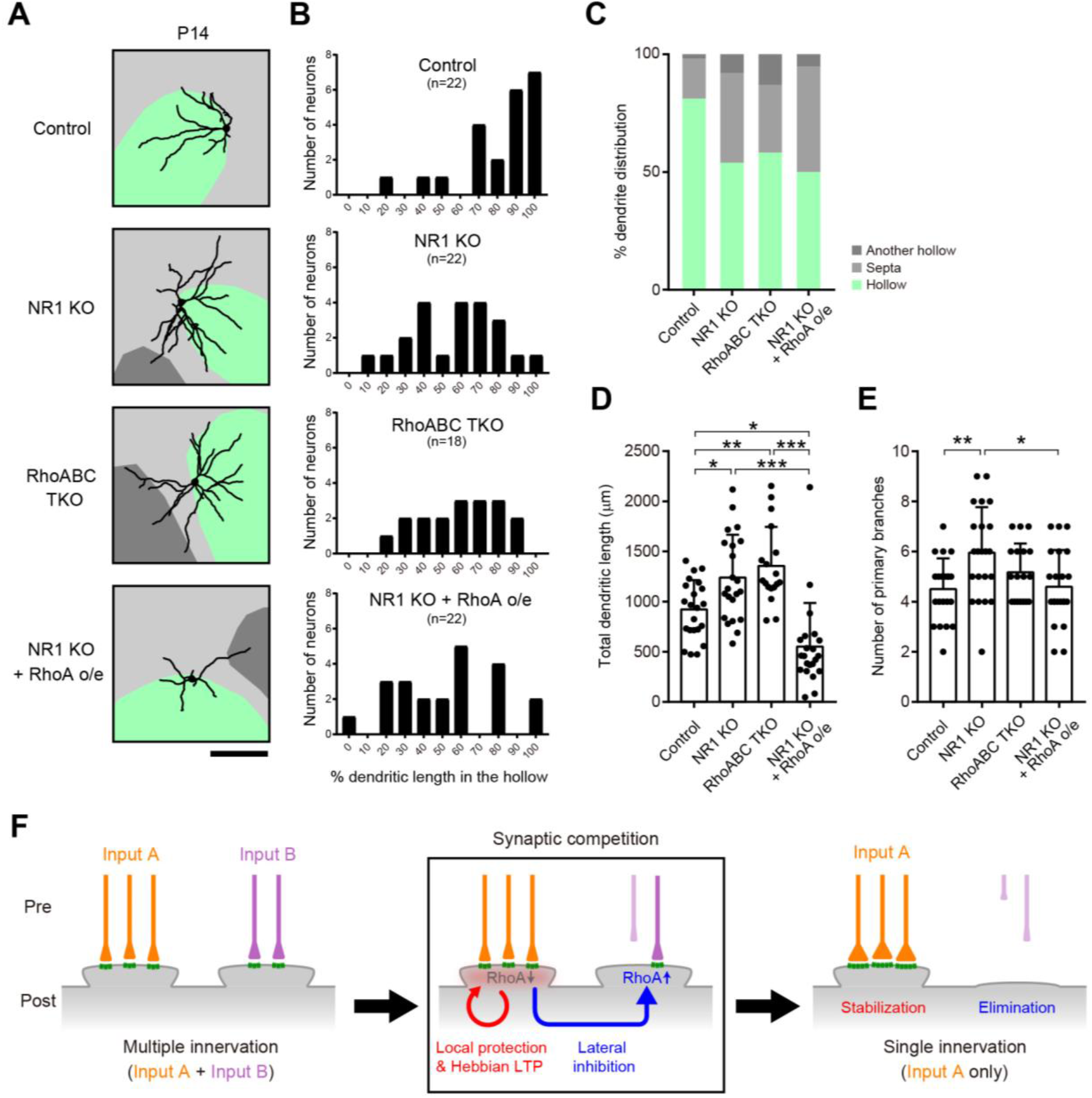
NMDAR-RhoA signals are essential for the synaptic competition in the barrel cortex. **(A)** Representative dendritic morphology of L4 neurons in the barrel cortex under control, NR1 KO, RhoA TKO, or NR1 KO + RhoA overexpression genotypes (*in utero* electroporation). Age, P14. Neurons in the barrel wall were analyzed. The somata of labeled L4 neurons located in the hollow shown in light green. Septa and neighboring hollows are indicated in light gray and dark gray, respectively. Hollow regions were determined by anti-VGluT2 immunostaining. The diameter of traced dendrites is thickened for visibility. Scale bar, 100μm. **(B)** Dendritic orientation. The *x*-axis of the histogram shows the percentage of the dendritic length within the major hollow (light green in **A**). **(C)** Quantifications of dendritic orientation. Mean ratios (%) of dendritic length in the major hollow, septa and adjacent hollows are shown. **(D)** Quantifications of total dendritic length. Data are the mean ± SD. *p<0.05, **p<0.01, ***p<0.001, (One-way ANOVA with post hoc Tukey’s multiple comparison tests). **(E)** Quantifications of the number of primary branches extending directly from the soma. Data are mean ± SD. *p<0.05, **p<0.01 (One-way ANOVA with post hoc Tukey’s multiple comparison tests). **(F)** A general principle of synaptic competition. Synapses receiving strong and correlated synaptic inputs are protected by RhoA suppression. They are also stabilized by Hebbian LTP. A subsequent neuron-wide activation of RhoA facilitates elimination of non-protected synapses, acting as the lateral inhibition signals.

## DISCUSSION

The circuit remodeling process is crucial for the establishment of discrete receptive field of a neuron. In this study, we found that glutamatergic inputs via NMDARs induce a local suppression as well as a neuron-wide activation of RhoA, facilitating lateral inhibition to leave just one type of synaptic inputs. In the olfactory system, the discrete connectivity of mitral cell dendrites to a single glomerulus is the anatomical basis for their discrete receptive fields in the odor space. Together with the “one neuron – one receptor” and the “one glomerulus – one receptor” rules for OSNs (Mori and Sakano, 2011), this discrete dendritic wiring, namely the “one mitral cell – one glomerulus” rule, ensures segregated olfactory information processing, from OSNs expressing a specific type of odorant receptor to distinct sets of mitral/tufted cells mediating specific odor perception and behaviors. Our finding is in stark contrast to the previous studies in the *Drosophila* olfactory system, in which dendrite wiring of projection neurons are defined by molecular cues (Hong and Luo, 2014).

The origins of spontaneous activity used in sensory circuit formation have been extensively studied in the visual, auditory, and somatosensory systems. For example, in the visual system, propagating spontaneous activity (known as retinal waves) generated in retinal ganglion cells regulates the fine tuning of the retinotopic map as well as eye-specific segregation in lateral geniculate nucleus (Feller *et al*., 1996; Huberman et al., 2008; Meister et al., 1991). In the auditory system, spontaneous ATP release activates hair cells, and subsequently afferent nerve fibers, contributing to tonotopic map formation (Babola *et al*., 2018; Tritsch *et al*., 2007; Wang et al., 2015). In the somatosensory system, the whisker-specific spontaneous activity is assumed to shape the whisker map in the somatosensory cortex (Anton-Bolanos et al., 2019; Mizuno *et al*., 2018). In this study, we showed that glutamatergic spontaneous activity generated within the olfactory bulb is critical for mitral cell remodeling. The pre-sensory spontaneous activity may be a general strategy to form sensory circuits before rich sensory experiences.

Activity-dependent synaptic competition is a widespread phenomenon in the developing nervous system. Earlier studies proposed the involvement of the “punishment signals” from winner synapses to the losers (Lichtman and Colman, 2000). However, the nature of the punishment signals has remained enigmatic. In this study, we showed that NMDAR-dependent activation of RhoA on a neuron-wide scale act as the “punishment signal” to prune loser dendrites (Figure 4-7). The global increase of RhoA is mediated by the depolarization of the neuron, at least in part. However, RhoA does not eliminate the winner dendrite, because NMDARs locally suppress RhoA to protect it from pruning (Figure 6H). In this way, regulation of RhoA implements activity-dependent lateral inhibition between dendrites. These mechanisms can now explain how a mitral cell establishes just one primary dendrite, not two or zero. A small difference in synaptic strength can be amplified after repeated glutamatergic inputs. If all dendrites accidentally retract from glomeruli, the dendrite elimination no longer proceeds without glutamatergic inputs, and dendritic regrowth might occur until at least one dendrite is connected to a glomerulus (Figure S7E and S7F).

Once one dendrite forms strong glutamatergic synapses, it will be further strengthened by Hebbian LTP via Rac1 (positive-feedback) (Aihara *et al*., 2021), while others will be further eliminated by lateral inhibition with RhoA (Figure 6H and 6I). Rac1 and RhoA seem to mediate actin fiber assembly and disassembly to facilitate neurite extension and retraction, respectively (Aihara *et al*., 2021; Luo, 2000). Notably, NMDARs are involved in synaptic competition in various systems (Personius et al., 2016; Rabacchi et al., 1992; Wong and Ghosh, 2002). Most likely, our proposed mechanism explains synaptic competition in other systems too (Figure 7F).

In activity-dependent circuit remodeling, synchrony is also known to be an important factor. Hebbian mechanisms are often used to explain this phenomenon, in which “Fire together, wire together” and “Out of sync, lose your link” control which connections remain (Hebb, 1949; Luo, 2021; Shatz, 1992). However, it has not been clearly explained why out-of-sync inputs are more effective for synapse elimination. In our current study using NMDA bath application, lateral inhibition worked efficiently when one dendrite is strong and the others are weak (Figure 6A, 6B, and 6E); however, it did not work when two dendrites receive sufficient amounts of NMDAR inputs simultaneously (Figure 6C, 6D, and 6F). We therefore assume that decorrelated inputs are useful to send lateral inhibition signals even when multiple dendrites already have strong glutamatergic synapses (Figure S7G). Over time, de-correlated mutual lateral inhibition could produce larger difference in synaptic strength. Thus, the decorrelated activity found at later stages of mitral cell development (Figure 2 and 3) may be helpful, if not essential, for an efficient dendrite pruning process. In the barrel cortex, multiple dendrites within the same hollow can be protected as they receive synchronized synaptic inputs during the remodeling process (Mizuno *et al*., 2018) (Figure 7 and Figure S7H).

Heterosynaptic plasticity is important not only during development, but also during the learning-related plasticity in various kinds of neurons (Chater and Goda, 2021; Chistiakova et al., 2014; Cichon and Gan, 2015). Synaptic competition is the most remarkable form of heterosynaptic plasticity. Lateral inhibition found in this study may be a general principle to form structured synaptic inputs and define discrete receptive fields of neurons, which are fundamental units of our brains. It will be interesting to study in the future how the different sources of signals, namely NMDAR-dependent local signals vs. depolarization signals, control opposite actions of RhoA.

## ACKNOWLEDGMENTS

We thank M. Yokoi (*Pcdh21-Cre*), I. Imayoshi (*R26-CAG-LoxP-TeNT*), P. Mombaerts (*OMP-GFP, OMP-Cre*), S. Tonegawa (floxed *Grin1, TRE-TeNT*), H. Zeng (*Ai38*), K. Svoboda (*Thy1-GCaMP6f*), C. Ron Yu (*OMP-tTA* knock-in), G.E. Shull (*NKCC1* KO) for mouse strains, C. Cepko (*pCALNL* and *pCAFNF* vectors), P. Soriano (*FLPo*), Y. Wu (DORA-RhoA), K. Aoki and M. Matsuda (RaichuEV-Rac1) for plasmids; A.S. Lowe for image processing software; I.D. Thompson for the neonatal anesthesia protocol; M. Okamoto and T. Miyata for instruction of *in utero* electroporation at E10; Y. Tsunekawa and F. Matsuzaki for instruction of CRISPR-Cas9; K. Aoki, H. Murakoshi, and R. Yasuda for valuable advices for slice FRET imaging; R. Iwata and M-T. Ke for valuable advices on imaging and analysis; T.R. Sato for comments on the manuscript. We are also grateful to M. Nomura, Y. Sakashita, Y. Taniyama, E. Yamashita, M. Nishihara, and T. Ohmine, and The Research Support Center, Research Center for Human Disease Modeling, Kyushu University Graduate School of Medical Sciences, for technical assistance. Imaging experiments were in part supported by the RIKEN Kobe Light Microscopy Facility and animal experiments were in part supported by the Laboratory for Animal Resources and Genetic Engineering at the RIKEN Center for Life Science Technologies. This work was supported by grants from the PRESTO and CREST (JPMJCR2021 to T.I.) programs of the Japan Science and Technology Agency (JST) (T.I.), AMED (JP20dm0207055 and JP21wm0525012 to TI), the JSPS KAKENHI (23680038, 15H05572, 15K14336, 16K14568, 16H06456, 17H06261, 21H00205, and 21H05696 to T.I., 15K14327, 17K14944, and 19K06886 to S.F., 17K14946 to M.N.L., JP18J10215 to S.A.), Mitsubishi Foundation (T.I.), Sumitomo Foundation (T.I.), Nakajima Foundation (T.I.), the Mochida Memorial Foundation for Medical and Pharmaceutical Research, the Uehara Memorial Foundation (T.I.), and RIKEN CDB intramural grant (T.I.). S.A. was a junior research associate at RIKEN and a predoctoral research fellow (DC2) of JSPS.

## AUTHOR CONTRIBUTIONS

S.F. performed *in utero* electroporation, morphological analysis, hand rearing, and Ca^2+^/FRET imaging of olfactory bulb slices. M.N.L. performed hand rearing and Ca^2+^ imaging of olfactory bulb *in vivo*. S.A. performed CRISPR-Cas9 screening and FRET imaging. R.S. performed *in utero* electroporation at E10. Y.M. and T.S. established *in utero* electroporation of mitral cells. R. K. and K.K. generated floxed *Cnga2*. T.I. supervised the project. S.F., M.N.L., S.A., and T.I. wrote the manuscript. S.F., M.N.L., and S.A. contributed equally to this work.

## DECLARATION OF INTERESTS

The authors declare no competing interests.

## STAR METHODS

### RESOURCE AVAILABILITY

#### Lead contact

Further information and requests for resources should be directed to and will be fulfilled by the Lead Contact, Takeshi Imai.

#### Materials availability

New plasmids generated in this study have been deposited to Addgene (https://www.addgene.org/browse/article/28197756/).

#### Data and code availability

Raw image data used in this study will be deposited to SSBD:repository (http://ssbd.qbic.riken.jp/set/2022xxxx/). Neurolucida tracing data will be deposited to NeuroMorpho.Org (http://neuromorpho.org/). Numerical data for all graphs are included in Table S2. Program codes have been deposited to GitHub (https://github.com/TakeshiImaiLab) as detailed below. Requests for additional program codes and data generated and/or analyzed during the current study should be directed to and will be fulfilled on reasonable request by the Lead Contact.

## EXPERIMENTAL MODEL AND SUBJECT DETAILS

### Mice

All animal experiments were approved by the Institutional Animal Care and Use Committee (IACUC) of the RIKEN Kobe Branch and Kyushu University. *OMP-tTA* BAC transgenic mice (*OMP-tTA* Tg; CDB0506T) and *R26-TRE-GCaMP3* BAC transgenic mice (*TRE-GCaMP3* Tg; CDB0505T) were generated in a previous study (Iwata *et al*., 2017). *Pcdh21-Cre* Tg (RBRC02189) (Nagai et al., 2005), *R26-CAG-LoxP-TeNT* knock-in (RBRC05154) (Sakamoto et al., 2014), floxed-*Cnga2* (Matsuo et al., 2015), floxed-*Grin1* (JAX #005246) (Tsien et al., 1996), *OMP-GFP* knock-in (JAX #006667) (Potter et al., 2001), *OMP-tTA* knock-in (JAX #17754) (Yu *et al*., 2004), *TRE-TeNT* Tg (JAX # 010713) (Nakashiba et al., 2008), *OMP-Cre* knock-in (JAX #006668) (Li et al., 2004), a Cre-dependent GCaMP3 knock-in line *Ai38* (JAX #014538) (Zariwala *et al*., 2012), *Thy1-GCaMP6f* Tg (line GP5.11) (JAX #024339) (Dana *et al*., 2014), *Nkcc1* KO (MMRRC #034262) (Flagella et al., 1999) have been described previously. ICR and C57BL/6N mice were provided from the Laboratory for Animal Resources and Genetic Engineering, RIKEN, or purchased from Japan SLC. ICR mice were used for *in utero* electroporation in Figures 1, 4, 5, 6, 7, S1, S2, S6 and S7. C57BL6N was used for Figure S1C. *Grin1*^*fl/fl*^, *OMP-Cre;Cnga2*^*fl/o*^, *OMP-Cre;R26-CAG-LoxP-TeNT, Pcdh21-Cre;R26-CAG-LoxP-TeNT, Pcdh21-Cre;Ai38* in C57BL/6 (C57BL/6N or C57BL/6J) or C57BL/6 × 129 (129P2 or 129S2) mixed background were bred to ICR mice for *in utero* electroporation (mixed at various ratios). *Nkcc1* KO backcrossed to FVB/N was bred to ICR mice for *in utero* electroporation (mixed at various ratios). Since the *Cnga2* gene is X-linked, male hemizygous mutant mice (*OMP-Cre;Cnga2*^*fl/o*^) were analyzed for *Cnga2*. For OSN-TeNT, we initially analyzed *OMP-tTA* knock-in mice (Yu *et al*., 2004) crossed with *TRE-TeNT* Tg (Nakashiba *et al*., 2008); however, TeNT was not expressed in all OSNs and a substantial amount of VAMP2 immunoreactivity remained in the OSN axon terminals (data not shown). We therefore used *OMP-Cre;R26-CAG-LoxP-TeNT* (both are knock-in), in this case VAMP2 immunoreactivity completely disappeared in OSN axons (Figure S2D). When mixed genetic backgrounds were used for quantification, samples from littermates were used for control experiments. Mice used for Ca^2+^ imaging (*OMP-tTA;TRE-GCaMP3, Pcdh21-Cre;Ai38, Thy1-GCaMP6f, Thy1-GCaMP6f;OMP-Cre;R26-CAG-LoxP-TeNT, Thy1-GCaMP6f;Pcdh21-Cre;R26-CAG-LoxP-TeNT*) were in C57BL/6 or C57BL/6 × 129 mixed background. Both males and females were used for our experiments. Mice were kept under a consistent 12-hour light/12-hour dark cycle (lights on at 8 a.m., and off at 8 p.m.).

## METHOD DETAILS

### Plasmid constructions

Cre-dependent pCALNL-GFP and Flp-dependent pCAFNF-DsRed are kind gifts from C. Cepko (addgene #13770 and #13771)(Matsuda and Cepko, 2007). *Kir2*.*1* (NM_008425.4) was PCR amplified from P10 mouse OB cDNA. Mutant *Kir2*.*1* (mutKir2.1)(Burrone *et al*., 2002) was generated using PCR-mediated mutagenesis. *TdTomato* (a gift from R. Tsien) was PCR-amplified and subcloned into pCALNL or pCAFNF vector. *Cre* (addgene #14797, a gift from C. Cepko), *iCre* (a gift from R. Sprengel), and *FLPo* (addgene #13793, a gift from P. Soriano) were PCR-amplified and subcloned into pCAG vector. FLAG-tag was fused at the C-termini of *Kir2*.*1* and *mutKir2*.*1*, and subcloned into pCALNL vector. *Dlg4* coding for PSD95 was PCR amplified from P10 mouse OB cDNA, and pCAFNF-PSDΔ1.2-GFP was generated as described previously (Arnold and Clapham, 1999; Hayashi-Takagi *et al*., 2015). We used PSDΔ1.2-GFP because the overexpression of full length PSD95 perturbed dendrite morphology in mitral cells (data not shown). Plasmids newly generated in this study will be deposited to Addgene (#125573-125581).

### *In utero* electroporation

For sparse fluorescent labeling, pCALNL-tdTomato (1 μg/μL) and pCAG-Cre (2-10 ng/μL) were used. To label mutant neurons sparsely, pCAFNF-tdTomato (1 μg/μL), pCAG-Flpo (2-10 ng/μL) were used. To generate single-cell *Grin1* KO mutants, pCAFNF-tdTomato (1 μg/μL), pCAG-Flpo (2-10 ng/μL), and pCAG-iCre (1 μg/μL) were used in homozygous floxed *Grin1* mice. *In utero* electroporation was performed as described previously (Saito, 2006). To label mitral cells at E12, 2 μL of plasmid solutions were injected into the lateral ventricle and electric pulses (a single 10-ms poration pulse at 72 V, followed by five 50-ms driving pulses at 32 V with 950-ms intervals) were delivered along the anterior-posterior axis of the brain with forceps-type electrodes (3-mm diameter, LF650P3, BEX) and a CUY21EX electroporator (BEX). *In utero* electroporation for E10 embryos was performed as reported previously (Okamoto et al., 2013). To label layer 4 neurons in the barrel cortex, *in utero* electroporation was performed at E13.5. All the details are summarized in Table S1.

### Naris occlusion

P0 mice were anesthetized on ice and a unilateral naris occlusion was quickly performed by cauterizing with soldering iron. Naris closure was confirmed under a stereomicroscope.

### Hand rearing

Hand rearing of newborn mice was performed as described previously (https://www.youtube.com/watch?v=sNX2byHbppM) with some modifications. Mice were kept in a boxed chamber (AsOne, #AS-600SC) maintained at 28-30°C and a humidity of 50%. A heater (Natsume, KN-475-3-40) and a water bath (Sansyo, SWS-181D) were used to maintain the temperature and humidity, respectively. Up to 3 pups were hand reared at the same time. Hand rearing was performed from P1 to P6. Pups were fed with Powdered Dog Milk at the recommended dilutions (Esbilac, PetAG, warmed to 37°C, ∼50μL/g) with a paint brush every ∼2 hours (done in shifts day and night). Their head and body were kept clean with a damp cotton bud. To facilitate digestion, excretion, and to prevent bloating, their abdomen was massaged with a cotton bud just after feeding. If the pups appeared dehydrated, a subcutaneous injection of saline was administered. A detailed step-by-step protocol will be published in Bio-protocols (Leiwe *et al*., 2020).

### Mouse brain samples

Mice were deeply anesthetized by an overdosing i.p. injection of pentobarbital (Dainippon Sumitomo Pharma). Anesthetized mice were perfused with 4% paraformaldehyde (PFA) in phosphate buffered saline (PBS). Excised brain samples were then fixed with 4% PFA in PBS at 4°C overnight. Samples were then embedded in 4% low-melting agarose (Thermo Fisher, #16520-100) to prepare brain slices. For immunohistochemical analysis, samples were cryoprotected with 30% sucrose at 4°C overnight and embedded in OCT compound (Sakura).

### Immunohistochemistry

Frozen OBs were cut into 16-μm-thick coronal sections using a cryostat (Leica). Sections were blocked with 5% normal donkey serum in PBS for 30 minutes and then incubated with primary antibodies in 5% normal donkey serum at 4°C overnight. After washing 3 times in PBS, sections were incubated with secondary antibodies for 1 hour at room temperature. Mouse anti-VAMP2 (1:500, Synaptic Systems, 104211) and mouse anti-FLAG (1:500, Sigma F3165) were used as primary antibodies. Alexa 555 or 647-conjugated donkey anti-mouse IgG (1:500, ThermoFisher, A-31570 or A-31571) are used as secondary antibodies. Nuclei were stained with 0.1% DAPI. Images were taken by a confocal microscope (Olympus, FV1000 or Leica SP8).

### Optical clearing and volumetric imaging

Mouse brain slices (2 mm-thick for SeeDB, 500 μm-thick for SeeDB2) were cut using a microslicer (Dosaka EM). Slices were then cleared with SeeDB (Ke et al., 2013; Ke and Imai, 2014) or SeeDB2 (Ke et al., 2016) as described previously. Step-by-step protocol and technical tips have been described in SeeDB Resources (https://sites.google.com/site/seedbresources/). SeeDB2 samples were stained with DAPI. Cleared tissues were mounted on glass slides with 2-mm or 500 μm silicone rubber spacer (Ke and Imai, 2014). Samples were imaged with an upright two-photon microscope (Olympus, FV1000MPE) with a motorized stage (Sigma-Koki). A commercialized 25x objective lens (Olympus, XLPLN25XSVMP, NA 1.0, WD 4.0 mm) were used to image SeeDB samples. InSight DeepSee Dual (Spectra-Physics) was used for two-photon excitation of tdTomato (1040 nm). SeeDB2 samples were imaged with inverted confocal microscopes: Olympus FV1000 with 405 nm, 488 nm, and 543 nm laser lines, or Leica SP8 with 405 nmand 552 nm laser lines.

### *In vivo* Ca^2+^ imaging of the olfactory bulb

#### Mouse lines

Transgenic *OMP-tTA;TRE-GCaMP3*(OSN-GCaMP3), *Pcdh21-Cre;Ai38*(M/T-GCaMP3), or *Thy1-GCaMP6f* (GP5.11) (M/T-GCaMP6f) were used for *in vivo* Ca^2+^ imaging. For the imaging of mutant mouse strains, they were crossed with the *Thy1-GCaMP6f* line (M/T-GCaMP6f). It should be noted that robust odor-evoked responses (ΔF/F > 100%) were detected in the adult olfactory bulb using these mouse lines (Iwata *et al*., 2017). Odor responses were also seen in the neonates (data not shown).

#### Anesthesia

Neonatal mice were anesthetized with age and weight specific doses and concentrations of ketamine/xylazine that were administrated by intra-peritoneal injections. 10mg/ml ketamine and 20mg/ml xylazine and H_2_O were mixed and injected as below.

Ratios (ketamine/xylazine/H_2_O in mL) and doses (/g body weight):

P0: 10:0.5:21, 0.025 mL/g; P2: 10:0.5:21, 0.02 mL/g; P4: 10:0.5:6, 0.015mL/g; P6-16: 10:0.5:6, 0.01 mL/g; P20-30: 10:0.5:6, 0.02 mL/g.

#### Surgery

Once anesthetized, a small craniotomy was made above the OB, and filled with Kwik-Sil (WPI). Then a 2mm circular coverslip (Matsunami) was placed above and sealed with superglue. Dental cement (Shofu) was then used to attach a custom-made bar (Narishige), allowing us to head fix with a custom head holding device (Narishige, MAG-2) and image once the mice had recovered.

#### Imaging

Post-surgery the mice were transferred to the head-fixed stage and placed under the microscope. The focal plane was set to the glomerular layer and then time-series imaging could begin. For the imaging of spontaneous activity, the pups were determined to be awake if they responded to a gentle tail pinch.

### Odor delivery

Amyl-acetate (Wako, 018-03623) was diluted in mineral oil (Sigma, M8410) (1 in 100) to a total volume of 1mL in a 50mL vial. The odor vapor was further diluted (1:10) in charcoal filtered air at a final rate of 2L/min. The air dilutions were performed with a custom-made olfactometer, at the following protocol 30 s baseline -> 5 s Odor -> 85 s rest for 5 trials. To synchronize imaging and the odor delivery TTL triggers were sent from the olfactometer to start imaging. All imaging was performed using the same water immersion objective lens (25x, Olympus, XLPLN25XWMP, NA=1.05, WD=2.0), at the same imaging settings (256 × 256, 0.429 seconds per frame).

### Ca^2+^ imaging of olfactory bulb slices *ex vivo*

*Pcdh21-Cre;Ai38*(M/T-GCaMP3), or *Thy1-GCaMP6f* (GP5.11) (M/T-GCaMP6f) were used for *ex vivo* Ca^2+^ imaging of olfactory bulb slices. Mouse pups were anesthetized on ice and sacrificed by decapitation. The olfactory bulb was quickly excised into ACSF (125 mM NaCl, 3 mM KCl, 1.25 mM NaH_2_PO_4_, 2 mM CaCl_2_, 1 mM MgCl_2_, 25 mM NaHCO_3_, and 25 mM glucose). OB slices were placed in a custom-made chamber. The chamber was placed under an upright two-photon microscope (Olympus, FV1000MPE) with a motorized stage (Sigma-Koki) and a water-immersion 25x objective lens (Olympus, XLPLN25XWMP, NA = 1.05, WD = 2.0). Before starting an imaging session, olfactory bulb slices are perfused with oxygenated ACSF (bubbled with 95% O_2_ and 5% CO_2_) at least for 1 hour at room temperature (25°C). During imaging sessions, olfactory bulb slices were continuously perfused with oxygenated ACSF (0.85-1.07 mL/min) at room temperature.

### Pharmacology

For Ca^2+^ imaging, 50 μM AP5 (Sigma, A5282), 25 μM CNQX (Sigma, C239), 300 μM Carbenoxolone (Sigma, C4790), 1mM 1-octanol (Sigma, 95446), 100 μM GABA (Sigma, A5835) and 1 μM TTX (Abcam, ab120055) were used. For FRET imaging, 100 μM NMDA (Nacalai, 22034-16), 40 μM Glycine (Sigma, G7126),100 μM AP5 and 1 μM TTX were used.

### Morphological analyses of mitral cell dendrites

Only the medial side of the olfactory bulb was analyzed. We only analyzed neurons located in the mitral cell layer. Therefore, external tufted cells and granule cells, sometimes labelled by *in utero* electroporation, were excluded from our analysis. We only quantified neurons where the entire dendritic field was included in the 3D image blocks. In our SeeDB protocol, glomeruli could be identified by autofluorescence signals. All primary dendrites were manually traced with Neurolucida software (MBF Bioscience). Feeding and quantification of dendrites were blinded in the hand rearing experiments. In all the remaining experiments, quantification was not blinded; however, to exclude any biases, all labelled neurons that meet the above criteria were quantified in an unbiased manner. Littermates were used as controls. Criteria for quantification are described in Figure 1H and 1I.

### Analysis of spontaneous *in vivo* Ca^2+^ imaging data

#### Motion and Drift Correction

Motion correction and drift correction were performed by using TurboReg (translation only, http://bigwww.epfl.ch/thevenaz/turboreg) then SPM (www.fil.ion.ucl.ac.uk/spm). Noise removal and subsequent ΔF/F movies were performed using AFID (a MATLAB plugin created by Dr. A. Lowe, https://github.com/mleiwe/AFID) (Lowe et al., 2013).

#### Spike Detection

Glomeruli were manually selected and ΔF/F values were calculated for each glomerulus. To detect glomerular spikes, a normal distribution was fitted to the lowest 80% of data points. Glomerular spikes were defined as points that were 3 standard deviations above the mean value calculated from a 10 second window around the time point in question.

#### Correlation Index

In order to measure the correlation of glomeruli a new measurement was created that measured the number of glomeruli active per frame (*G*^*n*^) as a percentage of the mean number of active glomeruli per frame 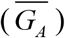. This normalization was necessary in order to control for differences in the number of glomeruli imaged as well as differences in the firing frequency of the network.

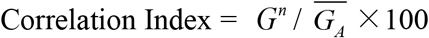

#### H- and L-events

H/L events were determined by calculating the number of glomeruli that were active in each frame. In Figures 2G, S3G, S4A, S6C and S6F, if the correlation index was above 5%, and contained two or more glomeruli it was classed as an H-event. Otherwise, if there were events present it was classed as an L-event

#### Spike Timing Tiling Coefficient (STTC) analysis

Calculations were performed as described previously (Cutts and Eglen, 2014). Correlations were counted within a 2 second window (i.e. ΔT=2 seconds).

Custom written MATLAB codes were used for all *in vivo* and *in vitro* analyses and are available at GitHub (https://github.com/mleiwe/SpontActivityOfDevMitralCells).

### Analysis of *in vivo* odor responses

#### Motion and Drift correction

Motion correction and drift correction was performed using TurboReg (translation only, http://bigwww.epfl.ch/thevenaz/turboreg), however with the shorter imaging times SPM was not necessary.

#### Glomerular Analysis

Glomeruli were manually selected and the ΔF/F values were calculated per glomerulus. F_0_ was defined as the mean value for the 15 seconds prior to odor application for each trial.

### Analysis of slice Ca^2+^ imaging data

Ca^2+^ imaging data were processed by using ImageJ (NIH) and MATLAB (Mathworks). Glomeruli were manually selected as polygonal region of interests (ROIs) on ImageJ. ROI information is converted to region mask using poly2mask function. Average fluorescence intensity within each ROI is defined as F. F_0_ is defined as the average of 10-30 percentile values of fluorescence intensity in each glomerulus. ΔF/F values were calculated for each glomerulus.

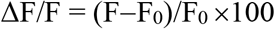

Motion correction and drift correction were not performed for *in vitro* imaging data analysis. Baselines were corrected by polynomial curve fitting. Raster plots were plotted by detecting peak points located in more than 4 standard deviations above the baselines. For calculation of standard deviations, top 25 percentile data were excluded. Correlation Index, H- and L-events were determined as described above. Cross correlation was calculated from ΔF/F. The value was normalized so that the autocorrelations at zero lag equal 1. *C*_*max*_ and *T*_*latency*_ were defined as shown in Figure S4D. Cross correlation matrix shows *C*_*max*_ values of all combinations of two glomeruli. Average frequency and amplitude were calculated from the number of peak points and ΔF/F value at each peak point, respectively.

### GCaMP images and movies

GCaMP fluorescence images shown in Figures 2, 3, 4, S2, S3 and S6 are maximum intensity projection images throughout the imaging session (also background subtracted) showing glomeruli. To prepare serial GCaMP images and ΔF/F images, **a** spatial median filter (diameter, 3 pixels) was applied to reduce the shot noise. For ΔF/F images, F_0_ values were created from the lowest 80% of pixel values. ΔF/F images are shown in pseudo-color (inferno). Unprocessed raw image data will be deposited to SSBD database (http://ssbd.qbic.riken.jp/set/2022xxxx/).

### FRET imaging of olfactory bulb slices *ex vivo*

pCAGGS-DORA-RhoA (van Unen et al., 2015) (a gift from Dr. Yi Wu) was modified as described previously (Murakoshi *et al*., 2011) to eliminate a cryptic poly A signal within the coding sequence (AATAA -> AACAA). Mutant DORA-RhoA has a mutation in PKN1 (L59Q) preventing RhoA binding (van Unen *et al*., 2015). pCAGGS-RaichuEV-Rac1 was described previously (Aihara *et al*., 2021; Komatsu et al., 2011). Plasmids were introduced at E12 by *in utero* electroporation. FRET imaging was performed at P3. Mouse pups were anesthetized on ice and sacrificed by decapitation. The OB was quickly excised into ACSF. Then OBs were embedded in 2% low melting point agarose and sliced into 300 μm-thick coronal sections using a microslicer (Dosaka EM). OB slices were put on a custom-made chamber and fixed by mesh (Warner Instrument). The chamber was placed under an upright two-photon microscope (Olympus, FV1000MPE) with a motorized stage (Sigma-Koki) and a water-immersion 25x objective lens (Olympus, XLPLN25XWMP, NA = 1.05, WD = 2.0). Before starting an imaging session, olfactory bulb slices are perfused with oxygenated ACSF at least for 2 hours at 27°C for recovery. During imaging sessions, OB slices were continuously perfused with oxygenated ACSF (0.85-1.07 mL/min) at room temperature (23-25°C). Excitation wavelength was at 840 nm. Emissions were split and filtered with FV10-MRC/YW filter cube (DM505, BA460-500 for CFP, and BA520-560 for YFP). Images were taken every 5-10 seconds (pixel size: 0.497 × 0.497 μm). In the single-cell analysis (Figures 6A-6D), multiple *z* planes were acquired (10-15 sections with 4-5 μm intervals) to cover multiple neurites. Other FRET images were acquired at single *z* planes.

### Analysis of FRET imaging data

FRET images were processed with ImageJ. Motion correction and drift correction were performed by Registration plugin (Correct 3D drift). Thresholding was performed based on the YFP channel using Auto Threshold function of ImageJ (Li or Huang method). Normalized YFP/CFP ratio was calculated, and ROI was manually drawn on all tufts and soma which were stably visible throughout experiments. In the data analysis, 2 min period before the stimulation was defined as a baseline and used for normalization. Averaged FRET signals during 2-4 min were quantified as responses in Figures 5C, 5F and 5G. Averaged FRET signals during 2-12 min were shown in single-cell analysis in Figures 6G. To generate representative FRET images, Gaussian filter (4×4 pixels) was used for Figure 5A, 5D, S7A and S7D and median filter (10×10 pixels) was used for Figures 6A and 6C.

### Single-cell knockout by plasmid-based CRISPR-Cas9 and guide RNA

pCAX-Cas9 and guide RNA backbone vector were kind gifts from F. Matsuzaki (Tsunekawa et al., 2016). For CRISPR-Cas9 based KO, three guide RNAs which target different exons were designed to increase KO efficiency (Aihara *et al*., 2021). Targeting sequences for *RhoA, RhoB, RhoC*, and *Grin1* are listed in Table S4. Target sequences were amplified with forward and reverse oligonucleotides by PCR and inserted into gRNA backbone vector at AflII sites (Tsunekawa et al., 2016). Guide RNA-resistant RhoA was generated by PCR-mediated mutagenesis and have the following sequences on the guide RNA-targeting sequences which code same amino acid sequences; GagAAtTAcGTaGCcGAcATaGAa (nucleotides, 118-141), AGtCCcGActcacTt (nucleotides, 262-276), and GAttTcaGaAAcGAtGAaCAtACccGA (nucleotides, 358-384) (small characters indicate modified nucleotides).

### Morphological analyses of L4 neurons in the barrel cortex

L4 neurons in the barrel cortex were labeled and manipulated by *in utero* electroporation at E13.5. Morphology was analyzed at at P14. After fixation, 120-μm-thick tangential vibratome sections of the primary somatosensory cortex were obtained. To visualize the barrel structures, whole-mount immunohistochemistry was performed as described previously (Ke *et al*., 2016). Brain sections were stained using guinea pig anti-VGluT2 (1:500, Millipore, AB2251-I) and Alexa 488-conjugated donkey anti-guinea pig IgG (1:500, ThrmoFisher, A-11073). Cell bodies and dendrites were reconstructed by Neurolucida software (MBF Bioscience). Neurons whose cell bodies are located on the barrel wall were analyzed. Spiny stellate and star pyramid neurons were not distinguished in the quantification. Note that dendrites sometimes extended outside the 120-μm-thick sections.

## QUANTIFICATION AND STATISTICAL ANALYSIS

Excel, Prism7, and MATLAB were used for statistical analysis. Sample sizes were not pre-determined. However, post-hoc power analyses determined that sufficient samples were collected. All the statistical data including the sample size and the statistical test used in each figure panel are described in Table S2. The number of neurons is described within figures, and all the numerical data including the numbers of animals are in Table S3. Number of animals used for Ca^2+^ imaging was at least three and indicated within figures. χ^2^-tests were used in Figures 1H, 1I, 1L, 1O, 4E, 4G, 4J, S1J, S1K, S2A, S2B, S5D and S6G-S6I. Multiple χ^2^-tests with Bonferroni correction were used in Figures 4H, 4I. Student’s t-tests (for parametric data), or Mann-Whitney U tests (for non-parametric data) were used in Figures 2G, 4C, 6G, S2I and S6C. Wilcoxon signed-rank tests were used in Figures 3D-3F, S5A and S5B. Kruskal-Wallis test and post hoc Dunn’s multiple comparison tests were used in Figure 1D. Welch’s t-tests were used in Figures S2C, S2E, S4A, S6B, S6C, S6F. Paired or unpaired t-tests were used for imaging data in Figures 5C, 5F,5G and S6E. One-way ANOVA and Tukey’s multiple comparison tests were used in Figure 5C, 7D and 7E. In box plots, the middle bands indicate the median; boxes indicate the first and third quartiles; and the whiskers indicate the minimum and maximum values. Data inclusion/exclusion criteria are described in figure legends.

## SUPPLEMENTAL FIGURES

**Figure S1.**
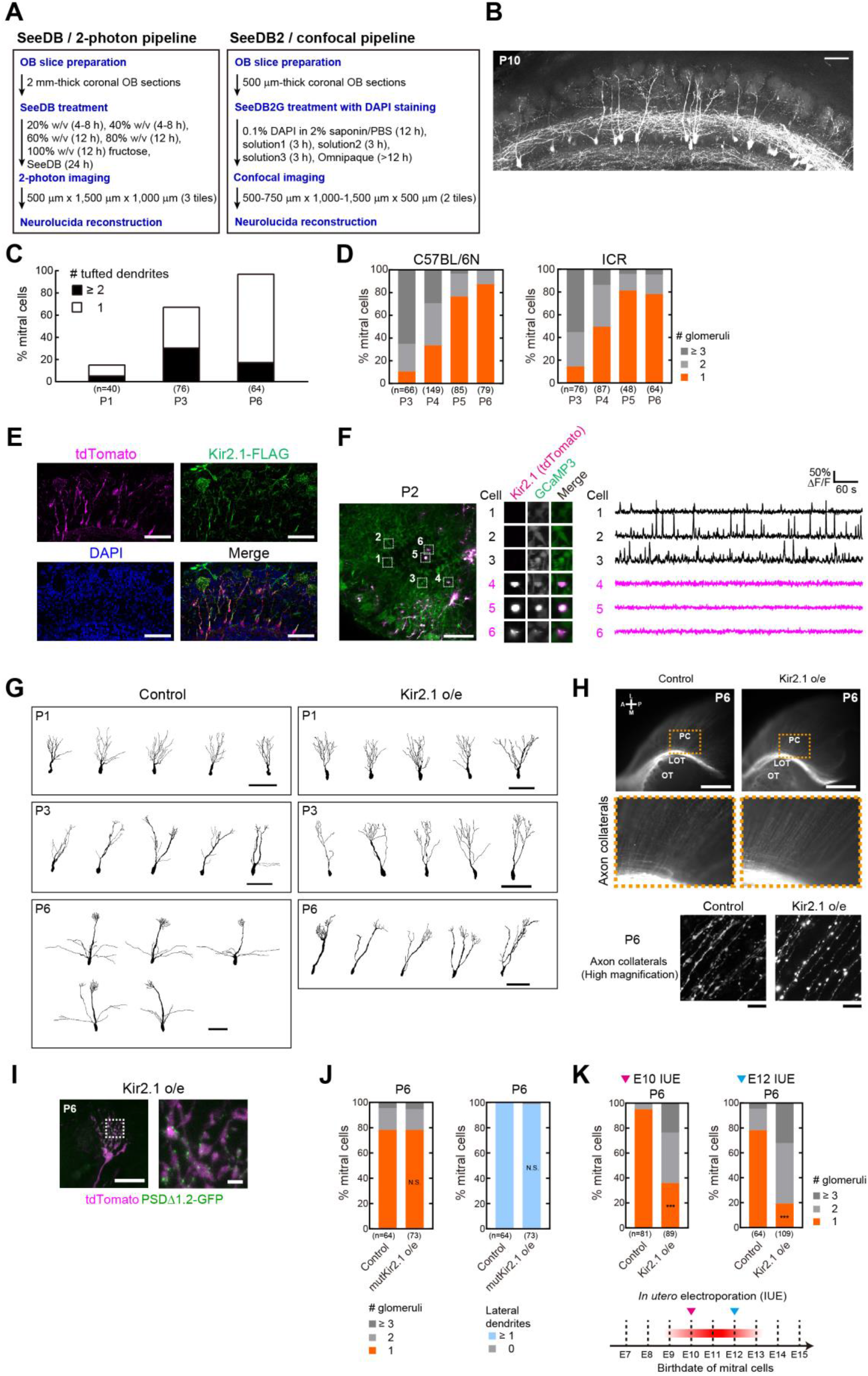
Quantitative analyses of dendrite pruning in mitral cells, related to Figures 1. **(A)** The clearing and imaging pipeline. OB slices were cleared with SeeDB or SeeDB2 and imaged using two-photon microscopy (excitation at 1040 nm for tdTomato) or confocal microscopy (excitation at 405nm for DAPI and 552 nm for tdTomato), respectively. **(B)** An example of mitral cells reconstructed by SeeDB-cleared olfactory bulb samples (100 μm stack) (see also Movie S1 for 1mm-thick images). The medial side of the OB was imaged by two-photon microscopy and all the labelled mitral cells were quantified in an unbiased manner. Autofluorescence signals were used to identify glomeruli for SeeDB samples. DAPI signals were used to identify glomeruli for SeeDB2 samples. A scale bar, 100 μm. **(C)** Quantitative analysis of the number of dendrites with a tufted structure at P1, P3 and P6. n, number of mitral cells. **(D)** Day-to-day quantifications of primary dendrites in C57BL/6N and ICR mice. n, number of mitral cells. **(E)** Immunostaining of FLAG-tagged Kir2.1. Almost all FLAG^+^ cells were positive for co-electroporated tdTomato (92.5%, 37/40 cells). Scale bars, 100 μm. **(F)** Spontaneous activity was not detected in Kir2.1-expressing mitral cells at P2 *ex vivo*. Kir2.1 was introduced to mitral cells in M/T-GCaMP3 mice using *in utero* electroporation at E12. A scale bar, 100 μm. Imaging was performed at 0.429 sec/frame. Kir2.1 was sparsely expressed together with tdTomato (left). The tdTomato-positive mitral cells had no spontaneous activity (right). A scale bar, 100 μm. **(G)** Representative control and Kir2.1-overexpressing mitral cells at P1, P3 and P6 reconstructed by Neurolucida. Scale bars, 100 μm. **(H)** Axonal projections of mitral cells expressing tdTomato alone (control, left) or with Kir2.1 (right). No obvious differences were seen in either the axonal extension in the lateral olfactory tract (LOT) and collateral formation in the piriform cortex (PC), at least at P6. OT, olfactory tubercle. Scale bars represent 1 mm. Fine scale images of axon collaterals are shown in the bottom. Scale bars represent 10 μm. **(I)** Postsynaptic structures on the primary dendrites in Kir2.1-overexpressing mitral cells at P6. Primary dendrites (magenta) and PSDΔ1.2-GFP puncta (green) are shown. Scale bars, 20 μm (left) and 2 μm (right). **(J)** Development of mitral cells expressing mutant Kir2.1 (mutKir2.1), a non-conducting mutant of the Kir2.1 potassium channel. Neither defects in the pruning process nor the lateral dendrite formation was observed (χ^2^-test vs control, N.S., non-significant), indicating that both defects by Kir2.1 overexpression are due to its hyperpolarizing effects. The control is the same as shown in Figures 1H and 1I. **(K)** Quantification of dendrite pruning for early-born mitral cells. *In utero* electroporation was performed for E10 embryos to label early-born mitral cells. The results were similar to E12-labelled ones (right, same data as shown in Figure. 1H).

**Figure S2.**
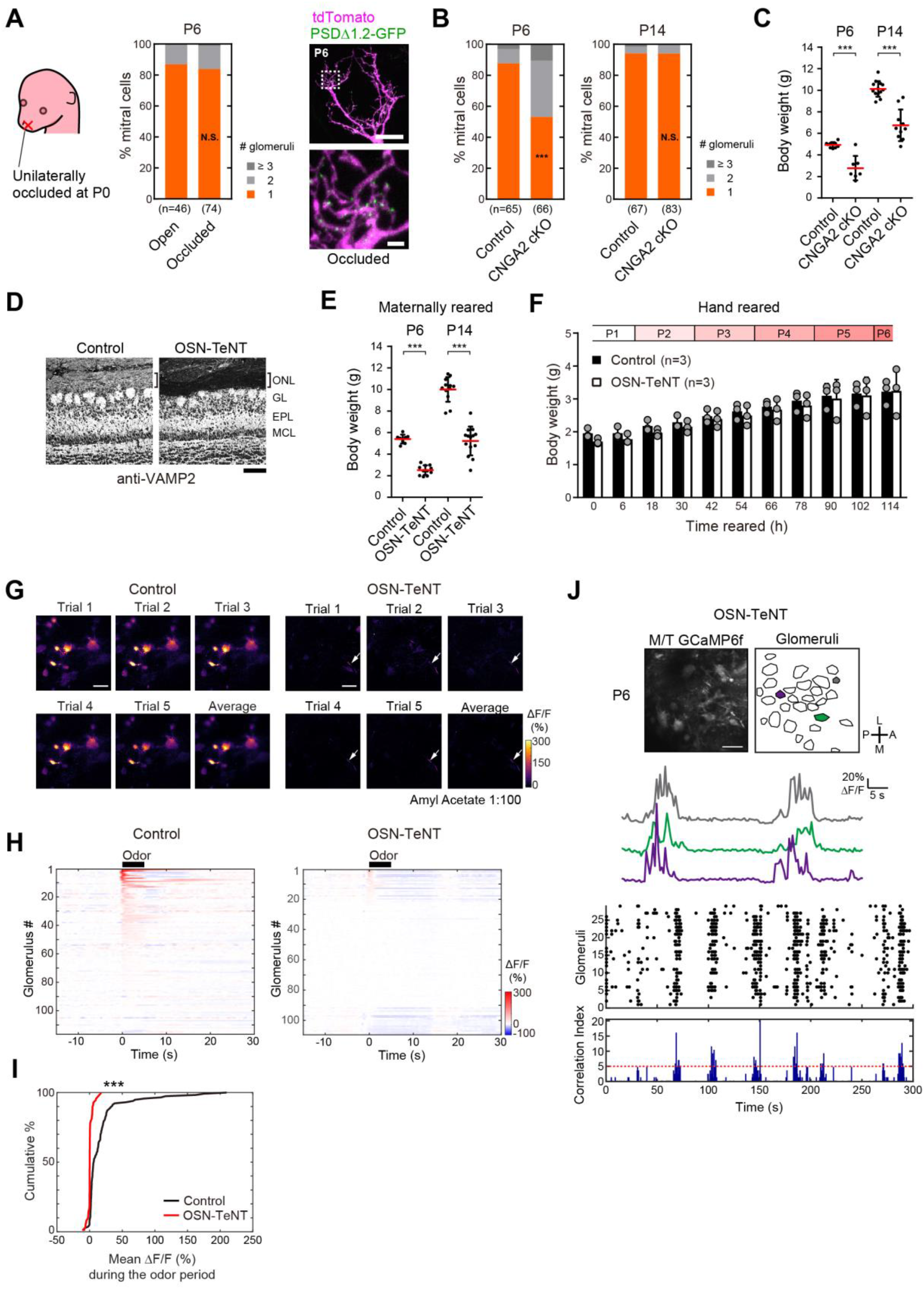
Manipulation of neuronal activity in OSNs, related to Figures 1. **(A)** Quantification of dendrite pruning after unilateral naris occlusion. N.S., non-significant (χ^2^ test, occluded vs open). Postsynaptic structures in naris-occluded mice are shown on the right. Scale bars, 20 μm (top) and 2 μm (bottom). **(B)** Quantification of dendrite pruning in OSN-specific CNGA2 mutant mice (*OMP-Cre;Cnga2*^*fl/o*^ male). ***p<0.001, N.S., non-significant (χ^2^ test vs control). **(C)** Quantification of body weight in control and CNGA2 cKO mice. CNGA2 cKO mice had significantly reduced their body weight. ***p<0.001 (Welch’s t-test). **(D)** OSN-TeNT mice *(OMP-Cre;R26-CAG-LoxP-TeNT*). VAMP2 immunostaining of olfactory bulb sections shows an almost complete depletion of VAMP2 signals in the olfactory nerve layer (ONL). GL, glomerular layer; EPL, external plexiform layer; MCL, mitral cell layer. A scale bar, 100 μm. **(E)** Quantification of body weight in maternally-reared control and OSN-TeNT mice. ***p<0.001 (Welch’s t-test). **(F)** Quantification of body weight in hand-reared control and OSN-TeNT mice. **(G)** Representative images of odor responses in control and OSN-TeNT mice showing that OSN-TeNT mice have significantly reduced odor responses. **(H)** Mean odor responses of glomeruli (5 trials) in control and OSN-TeNT mice (3 animals for both control and OSN-TeNT conditions). Black line indicates the delivery of amyl acetate at a 1:100 dilution. **(I)** Cumulative plot of the mean ΔF/F(%) value during the odor delivery period showing a marked reduction in odor responses, p<0.001, Mann-Whitney U test. **(J)** Imaging of maternally-reared OSN-TeNT mice crossed with M/T-GCaMP6f mice shows that spontaneous activity remains in mitral cells. Scale bars, 100 μm.

**Figure S3.**
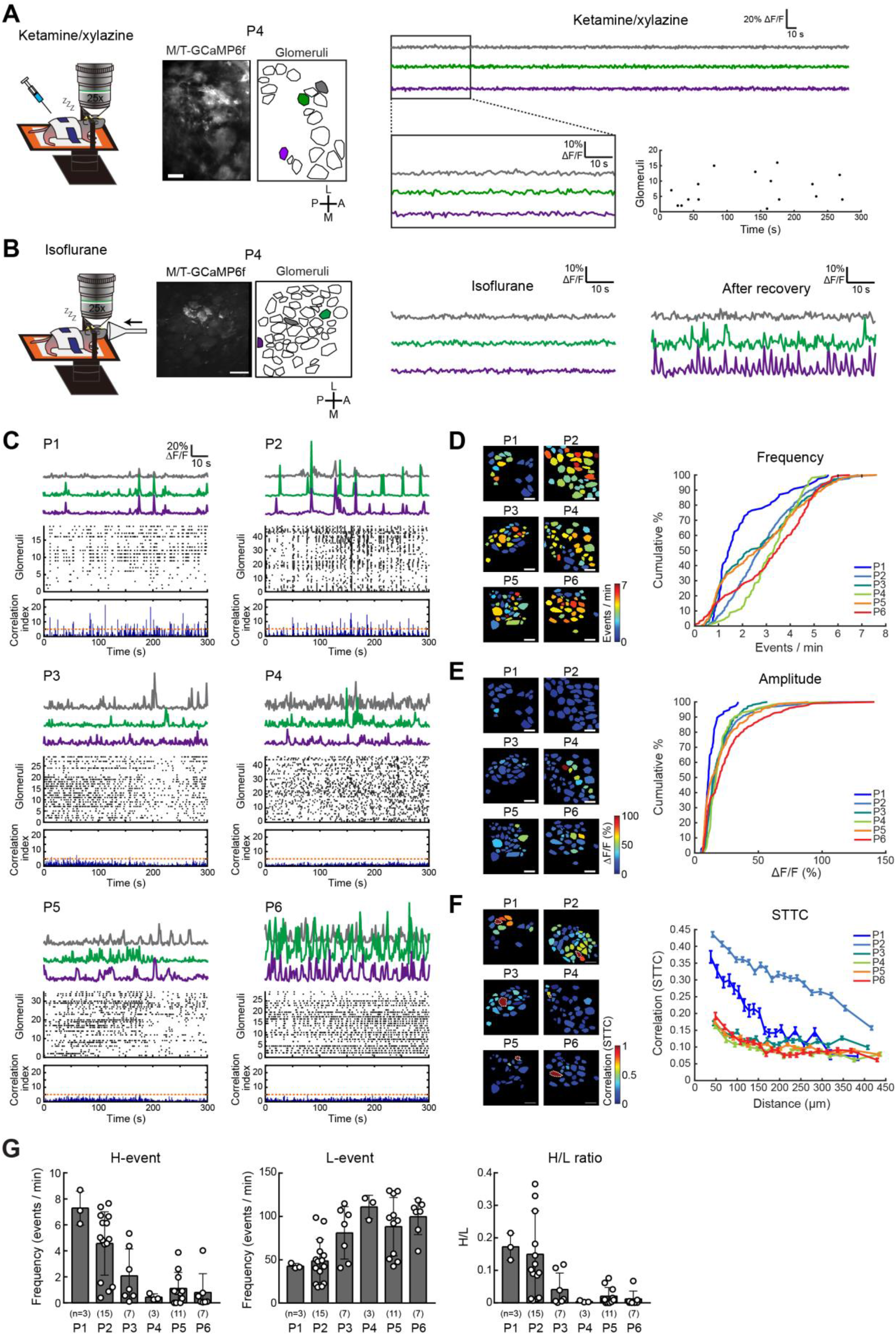
Developmental changes of spontaneous activity *in vivo*, related to Figure 2. **(A)** Schematic display of the ketamine/xylazine anaesthetized set up. Example traces for the 3 representative ROIs showing almost no spontaneous activity. Zoomed in area displays that the glomerular spikes are not evident. Raster plot displaying the spikes detected over a five-minute imaging period in the anesthetized mouse. **(B)** Example traces for the 3 representative ROIs under isoflurane anesthesia instead of ketamine/xylazine (left) and after recovery (right). Scale bars, 100 μm. **(C)** Representative traces, raster plots, and correlation index histograms (P1-6). Orange dotted lines indicate the threshold value for H- and L-events. P6 data is the same as in Figure 2. **(D)** Representative distributions of glomerular firing frequencies (P1-6, left). Cumulative distribution plots for the firing frequency of glomeruli (right). A data point represents the average firing frequency of a glomerulus over the imaging session. Data are for 77, 554, 210, 46, 394, and 222 glomeruli from 2, 5, 3, 3, 3, and 3 animals for ages P1-6, respectively. **(E)** Representative distributions of glomerular spike amplitude (P1-6, left). Cumulative distribution plots for the mean amplitude of glomerular spikes across all ages (right). A data point represents the average amplitude of spikes from a single glomerulus during an imaging session. **(F)** Spike Timing Tiling Coefficient (STTC) analysis shows the spatiotemporal relationship of the activity. P1 and P2 spontaneous activity showed a higher correlation within 100-150 μm, while later stages did not. Data points represent the mean ± SEM of glomerular comparisons into 20 groups, derived from 3, 15, 7, 3, 11, and 7 imaging sessions from 2, 5, 3, 3, 3, and 3 animals with total imaging times of 50.01, 309.95, 127.07, 57.87, 235.13, and 126.48 minutes for P1-P6 pups, respectively. Representative example heat maps for an example glomerulus (encircled by white dotted lines) are shown (right). Scale bars, 100 μm. **(G)** Developmental changes in H-event frequency, L-event frequency, and the H/L ratios (P1-6).

**Figure S4.**
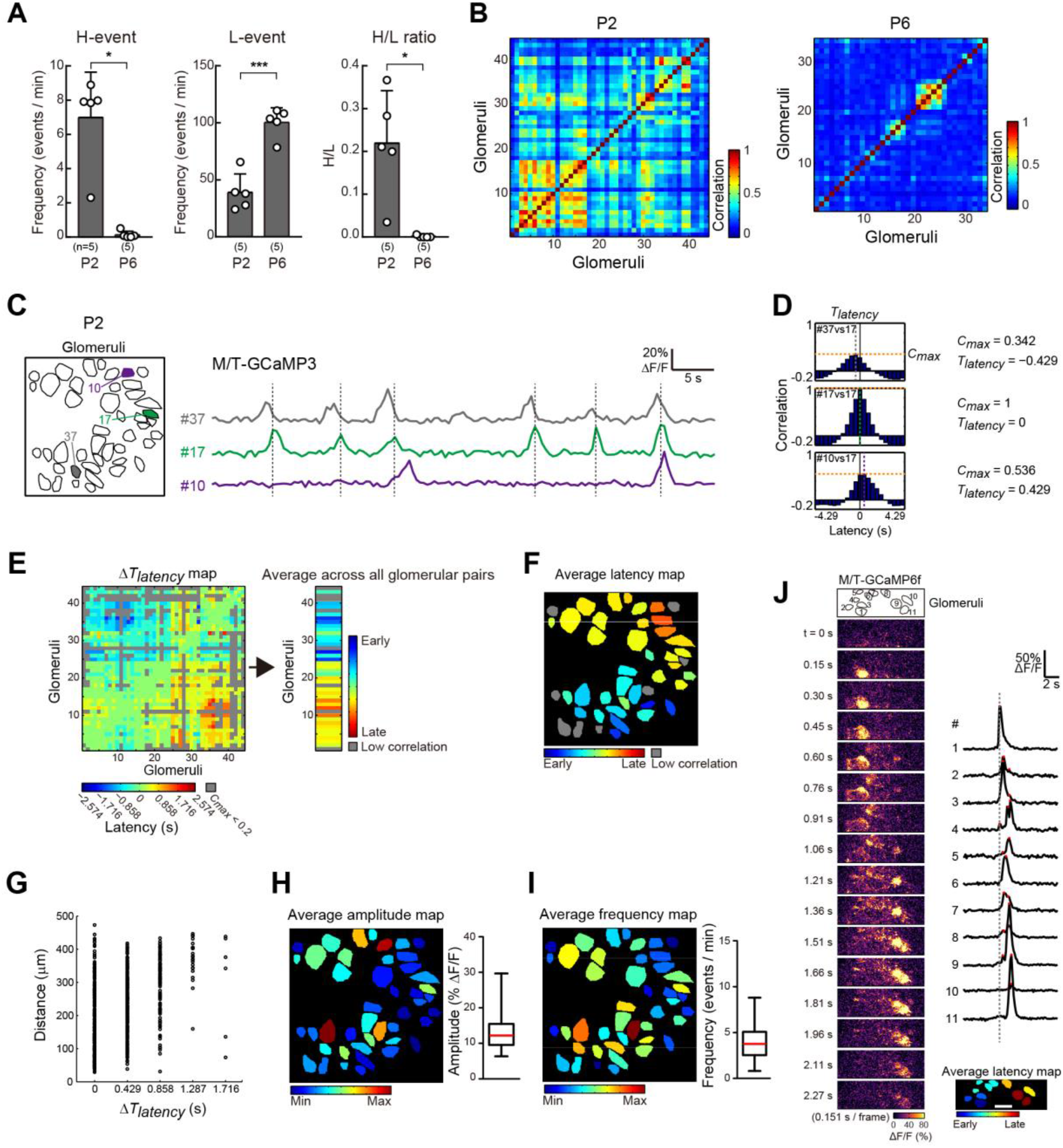
Spatiotemporal patterns of spontaneous activity at P2 and P6 *ex vivo*, related to Figure 3. **(A)** Developmental changes in H-event frequency, L-event frequency, and the H/L ratios in olfactory bulb slices. Data are mean ± SD. n=5 (P2), 5 (P6) sessions from 3 animals each. *p<0.05, **p<0.01 (Welch’s t test). **(B)** Cross-correlation matrices for P2 and P6 data. **(C, D)** Calculation of the maximum value of cross correlation (*C*_*max*_) and latency (*T*_*latency*_) to reach the *C*_*max*_. Raw traces **(C)** and cross correlograms **(D)** of 3 representative glomeruli are shown. **(E)** Δ*T*_*latency*_ map and average latency colormap. Glomerular pairs whose *C*_*max*_ values are smaller than 0.2 are shown in gray in the Δ*T*_*latency*_ map. Average latency was calculated based on the Δ*T*_*latency*_ map. Data were excluded from analysis when *C*_*max*_ values were <0.2 in more than half of glomerular pairs (shown in gray). **(F)** A latency map generated from the P2 olfactory bulb slice (shown in Figure 3B) demonstrates a propagating wave at this stage. The average latency was determined as in **(E)**. However, the average latency was mosaically arranged at a local scale. **(G)** The Δ*T*_*latency*_ and distance are plotted for all glomerular pairs. Glomerular pairs with *C*_max_ < 0.2 were excluded from this analysis. Based on this result, the propagation speed is estimated to be around 100-300 μm/sec *ex vivo*. Note that *ex vivo* experiments were performed at room temperature. **(H, I)** Average amplitudes **(H)** and frequencies **(I)** of glomerular spikes for P2 olfactory bulb slice (shown in Figure 3B). **(J)** High-speed Ca^2+^ imaging of olfactory bulb slices using M/T-GCaMP6f. Representative serial ΔF/F images of a propagating activity at P2 are shown. Ca^2+^ traces of individual glomeruli, show a propagating wave. Imaging was performed at 0.151 sec/frame. The average latency map is shown. A scale bar, 100 μm.

**Figure S5.**
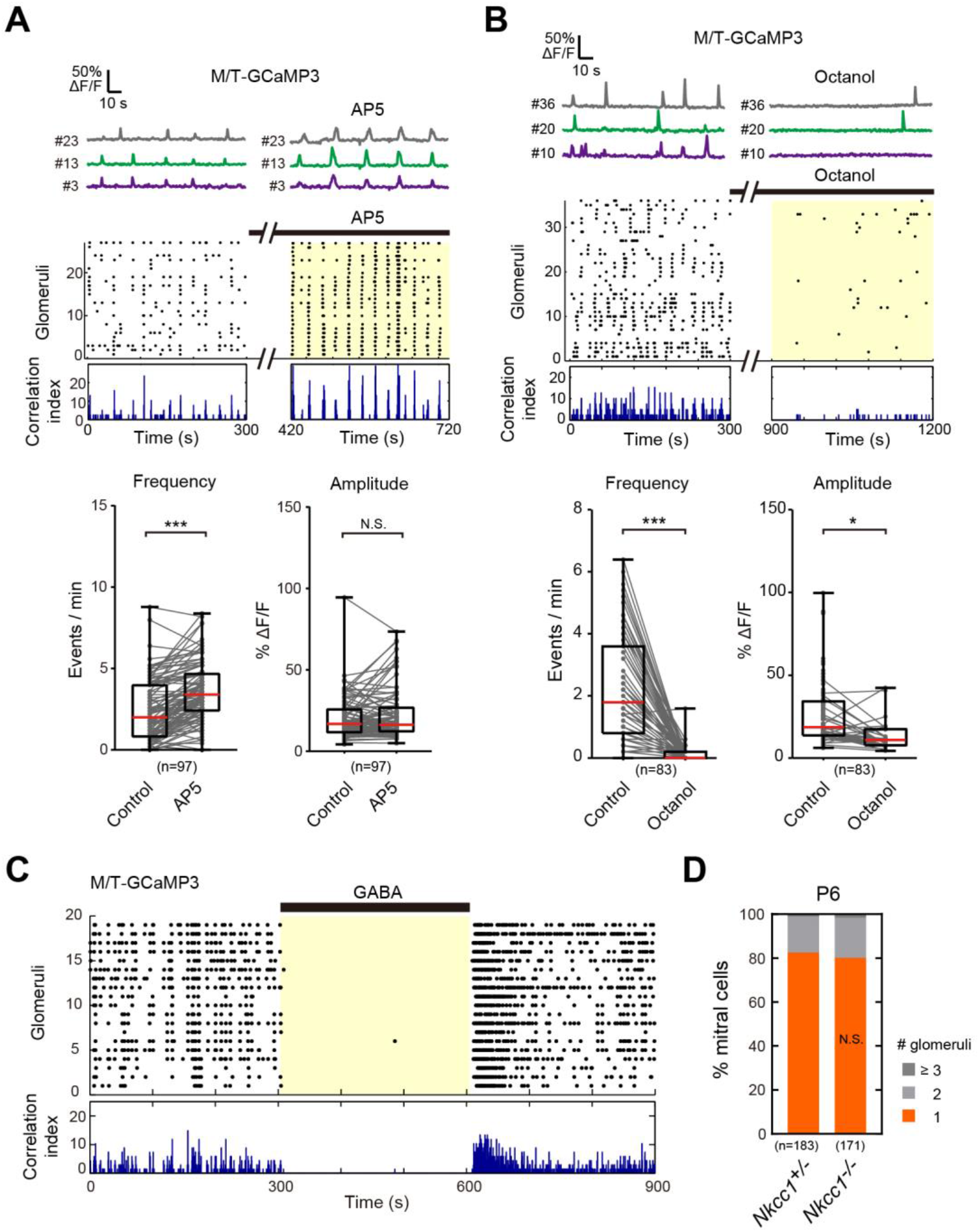
The role of NMDARs, gap junctions and GABA in generating spontaneous activity, related to Figure 3. **(A)** Spontaneous activity persisted when AP5 alone was added to P2-3 olfactory bulb slices. The pattern of activity was changed but the spontaneous activity remained. n = 97 glomeruli from 3 animals. N.S., non-significant, ***p<0.001 (Wilcoxon signed-rank test). **(B)** Ca^2+^ traces, raster plots, and correlation index histograms before and after Octanol (1 mM) application (P2) (left). Firing frequencies, but not amplitudes, were significantly reduced (right). n=83 glomeruli from 3 animals. *p<0.05, ***p<0.001 (Wilcoxon signed-rank test). **(C)** GABA (100 μM) was applied to P2 olfactory bulb slices. Spontaneous activity in mitral cells was almost completely suppressed by the application of GABA, suggesting an inhibitory role for GABA in mitral cells at this stage. **(D)** A knock-out mouse for NKCC1 did not show any defects in dendrite pruning in mitral cells. NKCC1 is a Na^+^-K^+^-2Cl^−^ cotransporter essential to incorporate Cl^−^ ion into cells and to make the neuron excitatory to GABA. N.S., non-significant (χ^2^-test compared to *Nkcc1*^*+/−*^).

**Figure S6.**
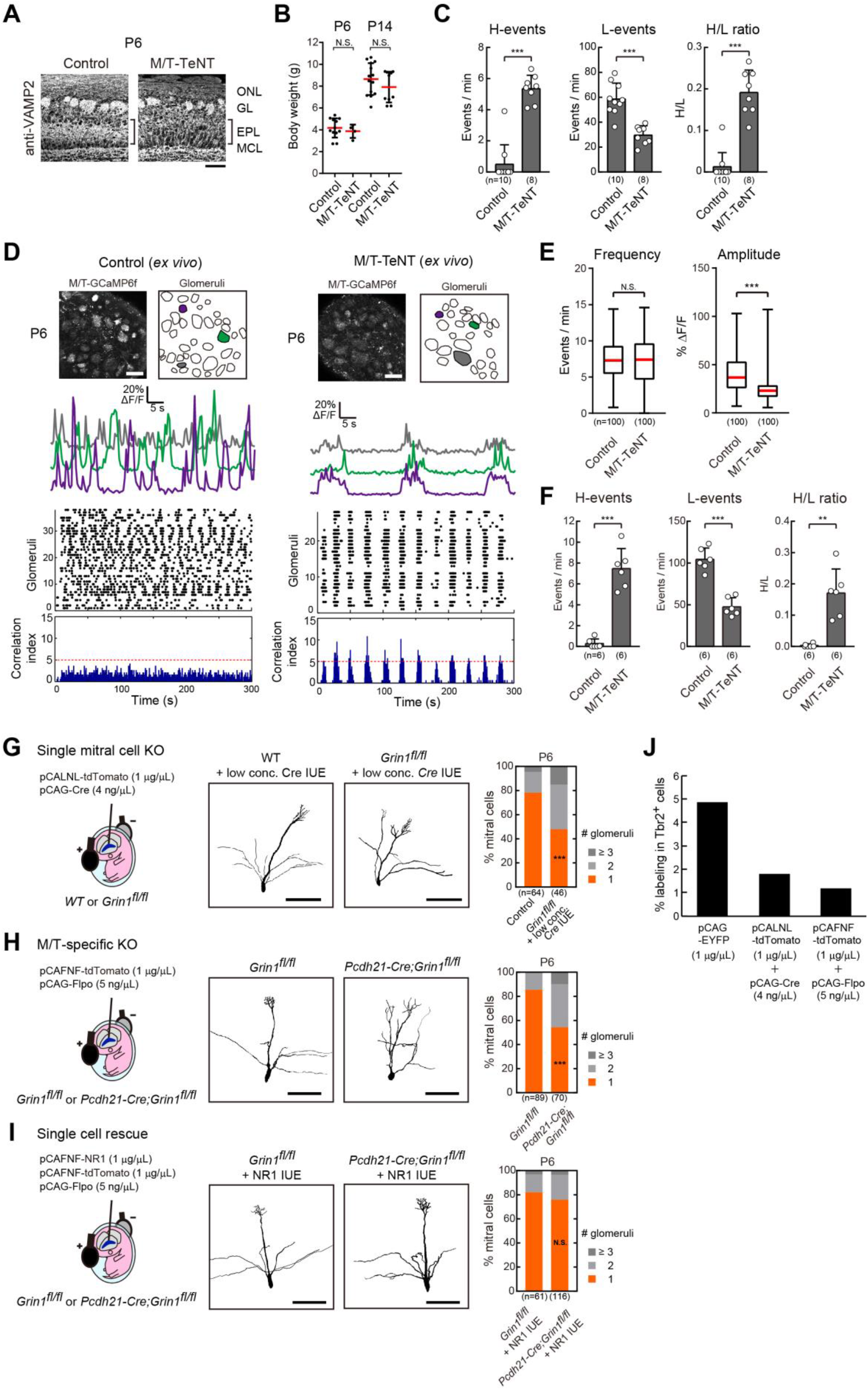
Genetic perturbation of glutamatergic neurotransmission in the OB, related to Figure 4. **(A)** M/T-TeNT mice *(Pcdh21-Cre;R26-CAG-LoxP-TeNT*). VAMP2 immunostaining was reduced in M/T-TeNT mice, particularly in the external plexiform layer of the olfactory bulb. A scale bar, 100 μm. **(B)** Quantification of body weight in control and M/T-TeNT mice at P6 and P14. N.S., non-significant (Welch’s t-test). **(C)** H-event frequency, L-event frequency, and the ratio of H to L events in control (littermates) and M/T-TeNT mice *in vivo*. ***p<0.001. As the H event frequency and H/L ratio were not normally distributed, Mann Whitney U tests were performed. For the L event frequency, a two-tailed t-test was performed. **(D)** Patterns of spontaneous activity in M/T-TeNT OB slices. Crossing M/T-TeNT mice with the M/T-GCaMP6f allowed us to visualize neuronal activity in M/T-TeNT mice at P6. Representative Ca^2+^ traces and their raster plots are shown. Scale bars, 100 μm. **(E)** The average frequency of glomerular spikes was comparable while the average amplitude was significantly decreased in the OB slices. N.S., non-significant, ***p<0.001 (Unpaired t-tests between control and M/T-TeNT). Data are from 100 glomeruli from 3 animals for both littermate controls and M/T-TeNT mice. **(F)** Measurements of H- and L-events show increased H/L ratio in M/T-TeNT OB slices. Data are mean ± SD. n=6 (control), 6 (M/T-TeNT) sessions from 3 animals each. *p<0.05, **p<0.01 (Welch’s t tests). Data are from 6 imaging sessions from 3 animals for both littermate controls and M/T-TeNT mice. **(G)** Sparse NR1 knockout with a low concentration of Cre (4 ng/μL) caused multiple primary dendrites. **(H)** M/T cell-specific NR1 knockout in *Pcdh21-Cre; Grin1*^*fl/fl*^ mice. **(I)** NR1 was sparsely expressed at a low concentration (5 ng/uL) of FLP and FLP-dependent NR1 plasmid in *Pcdh21-Cre; Grin1*^*fl/fl*^ mice. ***p<0.001, N.S., non-significant (χ^2^-tests vs. each control) in (G) - (I). Scale bars, 100 μm in (G) - (I). **(J)** Quantification of labeling densities with a high concentration of pCAG-EYFP (1 μg/μL) and low concentrations of Cre (4 ng/ μL) or FLPo (5 ng/ μL), combined with Cre/FLPo-dependent tdTomato. Only 4.87% of Tbr2^+^ cells were labeled with pCAG-EYFP (1 μg/μL). Low concentrations of Cre (4 ng/uL) and FLPo (5 ng/μL) labeled 1.81% and 1.17% of Tbr2^+^ cells.

**Figure S7.**
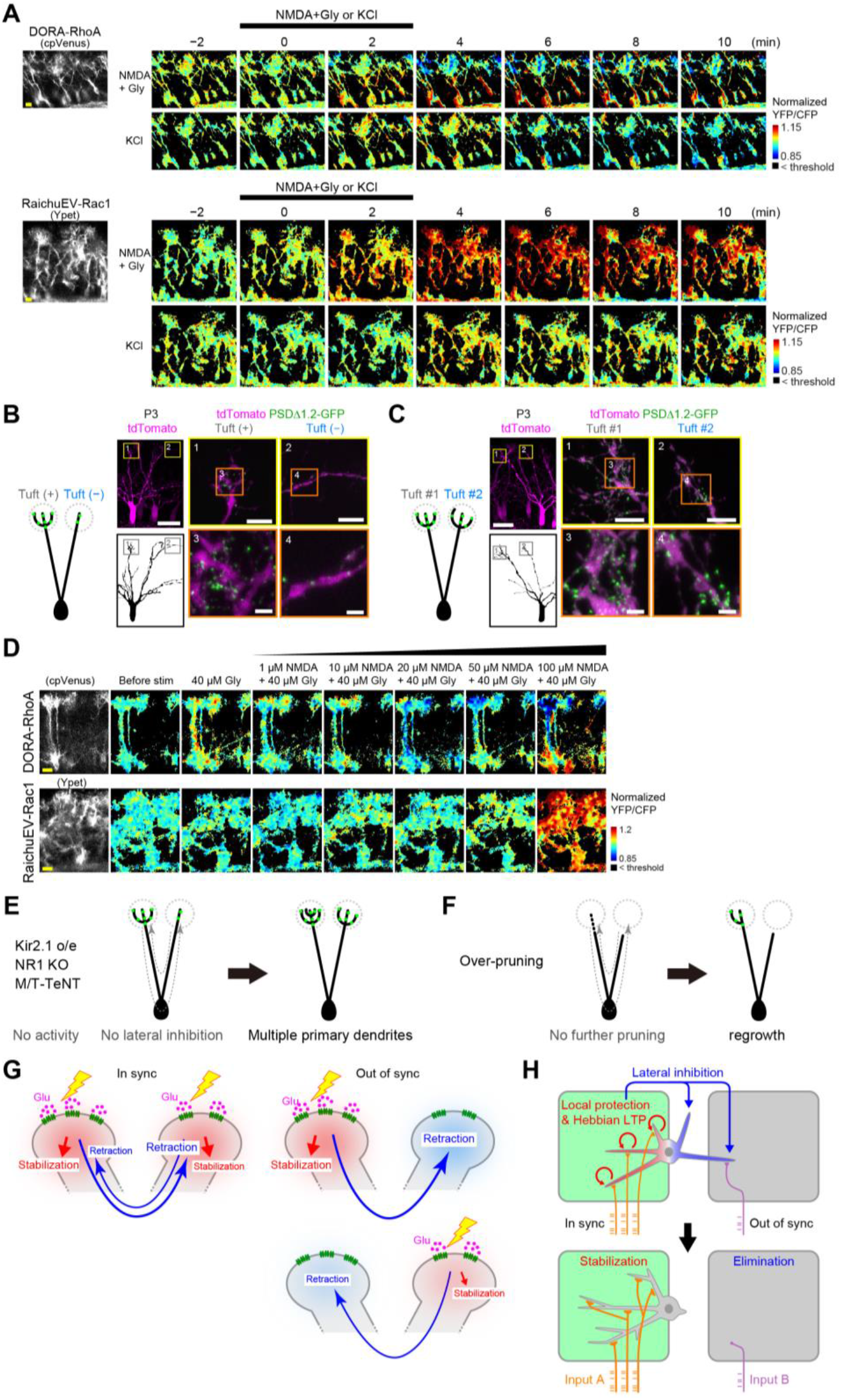
FRET imaging of Rho GTPases and models for dendrite pruning, related to Figure 5-7. **(A)** FRET imaging of RhoA and Rac1 activities with the DORA-RhoA and RaichuEV-Rac1 biosensors, respectively. Representative images of the tuft and soma regions are shown. RhoA and Rac1 samples are the same as in Figure 6A and 6D, respectively. **(B)** Distribution of glutamatergic synapses in a mitral cell with a tuft(+) and a tuft(−) primary dendrites. PDSΔ1.2-GFP (green) and tdTomato (magenta) were sparsely expressed in mitral cells. PSD puncta were mostly localized within the tufts. **(C)** Distribution of PSD puncta in mitral cells with two tuft(+) primary dendrites. **(D)** Representative images of dose responses for RhoA and Rac1 to NMDA. **(E)** When activity is blocked in mitral cells, there will be no pruning signals and multiple primary dendrites will be maintained. **(F)** Once a mitral cell accidentally loses all the connections from glomeruli, it will stop retraction without pruning signals, and would regrow the dendrite to form at least one connection. Thus, our model explains why a mitral cell acquire just one primary dendrite, but not two or zero. **(G)** Out-of-sync glutamatergic inputs may be more efficient for the lateral inhibition between dendrites. **(H)** A schematic model for the barrel circuit remodeling. Dendrites receiving the correlated inputs are all stabilized as they are protected from the lateral inhibition signals.

## SUPPLEMENTAL MOVIES

**Movie S1. Sparse labeling and optical clearing of OB**

Mitral cells were sparsely labelled with tdTomato using *in utero* electroporation and then the olfactory bulb was cleared with SeeDB. Glomeruli can be identified with autofluorescence. All the labelled neurons were quantified in an unbiased manner.

**Movie S2. Spontaneous activity *in vivo* at P2**

P2 olfactory bulb was imaged using two-photon microscopy. An awake M/T-GCaMP6f mouse was analyzed. Raw fluorescence (F) and ΔF/F images are shown.

**Movie S3. Spontaneous activity *in vivo* at P6**

P6 olfactory bulb was imaged using two-photon microscopy. An awake M/T-GCaMP6f mouse was analyzed. Raw fluorescence (F) and ΔF/F images are shown.

**Movie S4. Spontaneous activity *ex vivo* at P2**

P2 olfactory bulb was imaged *ex vivo* using two-photon microscopy. An M/T-GCaMP3 mouse was analyzed. Raw fluorescence (F) and ΔF/F images are shown.

**Movie S5. Spontaneous activity *ex vivo* at P6**

P6 olfactory bulb was imaged *ex vivo* using two-photon microscopy. An M/T-GCaMP3 mouse was analyzed. Raw fluorescence (F) and ΔF/F images are shown.

## SUPPLEMENTAL TABLES

**Table S1. IUE summary (Excel)**

Summary of IUE experiments.

**Table S2. Statistics data for figure panels (Excel)**

Statistics data for figure panels.

**Table S3. Numerical data for figure panels (Excel)**

Numerical data for figure panels. Each tab contains the numerical data for each figure panel.

**Table S4. gRNA target sequences (Excel)**

gRNA target sequences used for the CRISPR-Cas9 KO experiments.

## Notes

### Competing Interest Statement

The authors have declared no competing interest.

### Summary of Updates

In this revision, we showed that activity-dependent lateral inhibition via Rho GTPases establishes one primary dendrite per neuron. We also found that the NMDAR-RhoA signaling is essential for the synaptic competition in the barrel cortex.

